# Correlated activity favors synergistic processing in local cortical networks *in vitro* at synaptically-relevant timescales

**DOI:** 10.1101/809681

**Authors:** Samantha P. Sherrill, Nicholas M. Timme, John M. Beggs, Ehren L. Newman

## Abstract

Neural information processing is widely understood to depend on correlations in neuronal activity. However, whether correlation is favorable or not is contentious. Here, we sought to determine how correlated activity and information processing are related in cortical circuits. Using recordings of hundreds of spiking neurons in organotypic cultures of mouse neocortex, we asked whether mutual information between neurons that feed into a common third neuron increased synergistic information processing by the receiving neuron. We found that mutual information and synergistic processing were positively related at synaptic timescales (0.05-14 ms), where mutual information values were low. This effect was mediated by the increase in information transmission—of which synergistic processing is a component—that resulted as mutual information grew. However, at extrasynaptic windows (up to 3000 ms), where mutual information values were high, the relationship between mutual information and synergistic processing became negative. In this regime, greater mutual information resulted in a disproportionate increase in redundancy relative to information transmission. These results indicate that the emergence of synergistic processing from correlated activity differs according to timescale and correlation regime. In a low-correlation regime, synergistic processing increases with greater correlation, and in a high correlation regime, synergistic processing decreases with greater correlation.

**AUTHOR SUMMARY:** In the present work, we address the question of whether correlated activity in functional networks of cortical circuits supports neural computation. To do so, we combined network analysis with information theoretic tools to analyze the spiking activity of hundreds of neurons recorded from organotypic cultures of mouse somatosensory cortex. We found that, at timescales most relevant to direct neuronal communication, neurons with more correlated activity predicted greater computation, suggesting that correlated activity does support computation in cortical circuits. Importantly, this result reversed at timescales less relevant to direct neuronal communication, where even greater correlated activity predicted decreased computation. Thus, the relationship between correlated activity and computation depends on the timescale and the degree of correlation in neuronal interactions.

---

What role does the correlated activity among cortical neurons play in neural information processing? Correlated activity is ubiquitous throughout the brain, emerging from both external stimuli and internal dynamics. Correlated activity is predictive of information processing (for a review, see Salinas & Sejnowski, 2001). However, the extent to which it is favorable for information processing is not clear. What is needed to better understand the role of neural correlations in information processing is a comparison of how the amount of correlated activity between upstream neurons relates to the amount of resulting information processing in cortical microcircuits.

The view that correlated neural activity is favorable for neural information processing is widely held within the cognitive rhythms community and is based on the idea that correlation facilitates both communication between circuits and the orchestration of processing within circuits. Correlated neural activity, especially synchronous activity, is understood to generate the coherent rhythms that are observed in local field potentials, electrocorticography, and electroencephalography which are theorized to subserve specific computational or cognitive mechanisms (e.g., Fries, 2015; Hasselmo et al., 2002; Honey et al., 2017; Lisman & Jensen, 2013; Newman et al., 2014; Norman et al., 2006; Ward, 2003; Hernandez et al., 2020). Ample empirical evidence derived from *in vitro*, *in silico*, and *in vivo* studies supports the importance of synchrony for organizing information transmission in cortical circuits (Averbeck & Lee, 2004; Azouz & Gray, 2003; Fries, 2015; Poulet & Petersen, 2008; Salinas & Sejnowski, 2001; Yu et al., 2008). The synchronization of neuronal spiking is indeed linked to higher order cognitive and behavioral processes (Grammont & Riehle, 1999; Riehle et al.,1997; Vinck et al., 2015). From this perspective, correlation is favorable for information processing. Yet, even within the cognitive rhythms community there is recognition that excess correlation can be unfavorable and that, in some circumstances, desynchronization supports information processing better than synchronization (e.g., Bastos et al., 2015; Hanslmayr et al., 2012; van Winsun et al., 1984). The discrepancy between this and the standard view of the cognitive rhythms community warrants reconciliation (Hanslmayr, Staresina, & Bowman, 2016).

The view that correlated activity is unfavorable for neural information processing is widely held in the sensory processing and artificial neural network communities and is based on the idea that correlation is synonymous with redundancy and thus reduces efficiency (Attneave, 1954; Barlow, 1961; Gutnisky and Dragoi, 2008; Schneidman, Bialek & Berry, 2003; Shadlen & Newsome 1998). This bandwidth-limiting effect of correlation motivates the ‘redundancy-reduction hypothesis’ (Atick & Redlich, 1990; Attneave, 1954; Barlow, 1961; Shadlen & Newsome 1998). In this line of thinking, a normative goal of sensory information processing is to reduce the redundancy of neuronal signals (Atick and Redlich, 1992; Barlow, 1961; Field, 1987; Gutnisky and Dragoi, 2008; Laughlin, 1989; Rieke et al., 1995; van Hateren, 1992). Many works, together, indicate that signal redundancy decreases from lower order sensory areas to higher order sensory areas (Berry et al., 1997; Chechik et al., 2006; Dan et al., 1998; Doi et al., 2012; Montani et al., 2007; Nirenberg et al., 2001; Puchalla et al., 2005; Reich et al., 2001; Reinagel and Reid, 2000; Zohary et al. 1994). From this perspective, correlation is unfavorable for information processing. Yet, as in the cognitive rhythms community, there is an indication that qualification is needed to this standard view, given recent empirical evidence that correlated activity can also increase processing (Nigam, Pojoga, & Dragoi, 2019).

Our aim with the work described here was to determine which of these two perspectives better accounts for the relationship between correlation and synergistic processing—a component of information processing—in local cortical microcircuits at synaptic timescales and beyond. To accomplish this goal, we analyzed the spiking activity of hundreds of neurons recorded simultaneously from each of 25 organotypic cultures of mouse somatosensory cortex (Fig 1). In these recordings we identified hundreds of thousands of triads wherein individual neurons received significant functional input from two other neurons at synaptic timescales (< 14ms). For each triad, we measured the total information about the receiving neurons firing that was carried in the sending neurons activity and then decomposed this into the constituent components: the unique contributions of each sender, the redundancy between the senders (i.e., redundancy), and the synergy between the senders (i.e., synergistic processing). Across triads, we found that the correlation between the senders was low but that greater correlations were predictive of greater synergistic processing by the receiving neuron. When we extended the timescale of the analysis to consider triads at extrasynaptic windows (up to 3000ms), we observed a significant increase in the overall correlation between unique senders’ activity. We also observed a shift to a negative relationship between sender correlation and synergistic processing. Secondary analyses of the redundancy and information transmission indicated that both perspectives are valid but become relevant in different correlation regimes. Specifically, in low-correlation regimes (at synaptic timescales), information transmission and redundancy also increased, but the rate of increase of information transmission was greater than redundancy, thereby enabling greater synergistic processing. In high-correlation regimes (at extrasynaptic timescales), information transmission leveled off but redundancy continued to increase, thereby consuming bandwidth capacity and reducing synergistic processing.

**Figure 1.**
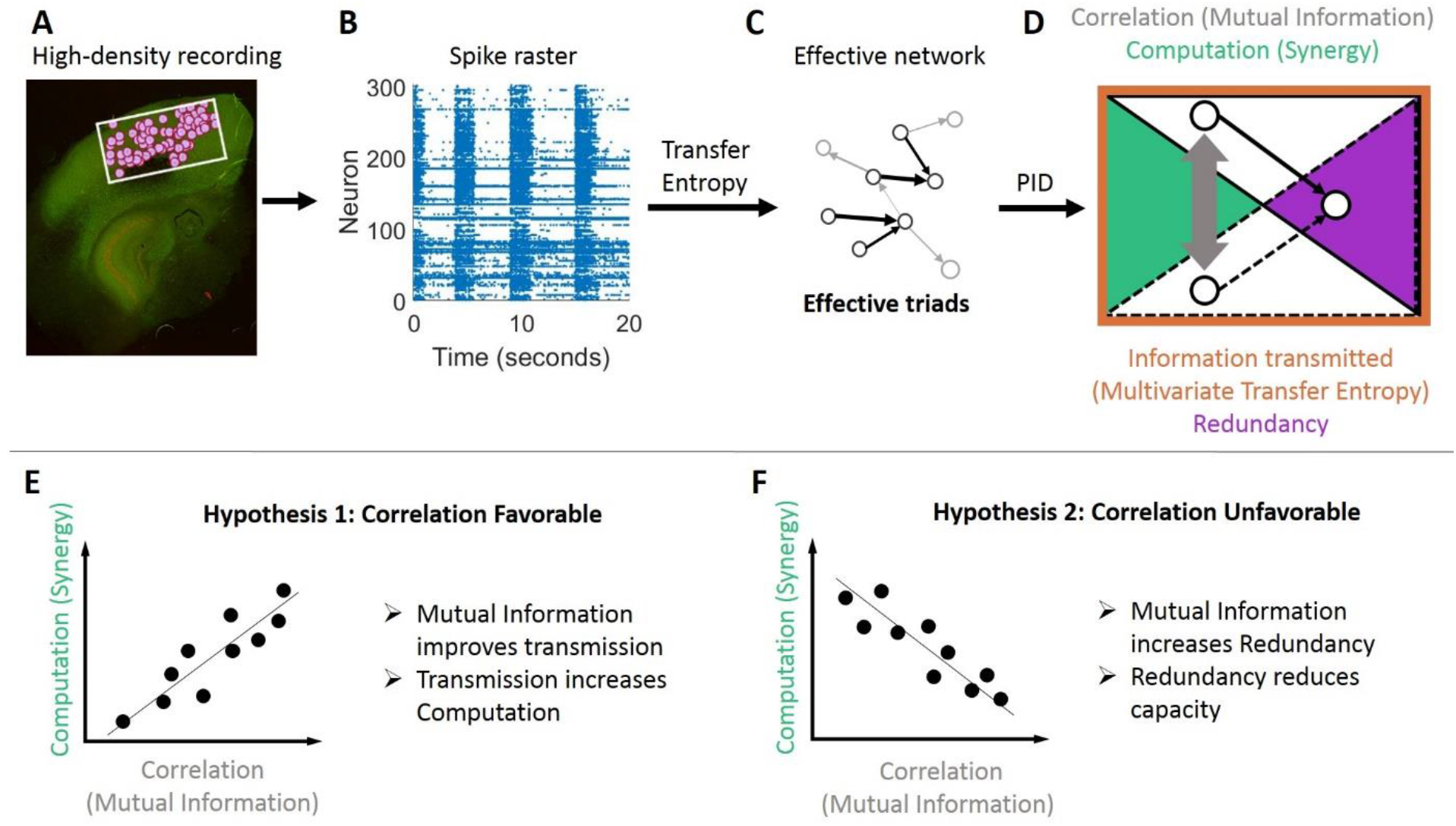
Methodological approach taken to ask if correlated activity is favorable or unfavorable for synergistic processing in organotypic cultures of mouse cortex. (A) Hour-long recordings of spiking activity were collected *in vitro* from organotypic cultures of mouse somatosensory cortex. (B) Spike sorting yielded spike trains of hundreds of well-isolated individual neurons. (C) Effective connectivity between neurons was determined by quantifying transfer entropy between spike trains. The resulting effective networks were analyzed to identify all effective triads consisting of two effective connections to a common receiver. (D) For each triad, we quantified how correlated the activity of the senders was with mutual information (gray arrow) and decomposed the total information transmitted to the receiver (measured via multivariate transfer entropy, orange perimeter) into redundancy and synergy components (via partial information decomposition, indicated with purple and green areas respectively). (E-F) We sought to differentiate between two alternate hypotheses. (E) Predicted relationship between mutual information and synergy if correlated activity is favorable for computation. (F) Predicted relationship between mutual information and synergy if correlated activity is unfavorable for computation.

## RESULTS

We asked whether correlated activity between neurons supports computation in cortical microcircuits by analyzing hour long recordings of spiking activity from organotypic cultures of mouse somatosensory cortex (n = 25) as summarized in Figure 1. Recordings contained between 98 and 594 well-isolated neurons (median = 310). We identified effective connections between neurons in each recording as those that had significant transfer entropy. For every effective triad consisting of one neuron receiving effective connections from two other neurons we quantified the amount of correlated activity between the sending neurons using mutual information. Mutual information was selected for its ability to detect nonlinear as well as linear dependencies. Our findings were, however, not sensitive to the method for quantifying correlated activity (see Supplemental Materials where we show the same qualitative pattern of results using Pearson correlation as well as other measures). The mutual information between senders was normalized to reflect the proportion of the maximum possible mutual information for that triad given the constituent neuron entropies (*p*MI^max^). We quantified the amount of computation performed by the receiver on the inputs using ‘synergy,’ a term derived from partial information decomposition. Synergy was normalized to reflect the proportion of the receiving neuron entropy for which it accounted (*p*H^rec^). Neither normalization was required (as shown in the Supplemental materials) but were included to control for variability across networks. Across triads, we asked whether synergy was positively or negatively correlated with mutual information. This analysis was repeated at varying timescales as determined by the granularity of the data binning and the delay between bins. All summary statistics are reported as medians followed by 95% bootstrap confidence intervals in brackets.

### Correlated activity is associated with more synergy at synaptic timescales

To determine the relationship between mutual information and synergy at timescales relevant for synaptic processing, we analyzed the functional dynamics at three timescales spanning 0.05 - 14 ms (centered on 3, 5, and 11 ms) for each of the 25 recordings, yielding 75 total functional networks. Each network had many effective triads (count = 806 [448 1350]). The median mutual information across triads of individual networks was 0.008 [0.005 0.010], which was ~0.8% of the maximum possible mutual information. Mutual information was highly variable around this median value within individual networks, varying across ~7.0 [6.7 7.2] orders of magnitude within individual networks. We then asked how this variability in mutual information was correlated with variability in synergy.

Plotting synergy as a function of mutual information across triads for each network revealed clear positive relationships (e.g., Fig 2A). To aggregate results across networks with differing numbers of triads and to maintain the ability to observe possible non-monotonic relationships between mutual information and synergy, we collapsed results across triads based on mutual information deciles for each network (Fig 2B). Mutual information and synergy were strongly positively correlated across deciles (Spearman *r* = 0.95 [0.93, 0.98]; Z_s.r._ = 7.53, n=75 networks, p <1×10^−13^; Fig 2C). The median slope over deciles in the log-log space was 0.46 [0.41 0.50] Δlog_10_(*p*H^rec^) / Δlog_10_(*p*MI^max^) (Fig 2D) indicating that synergy increases exponentially as a function of mutual information with an exponent of ~0.46. This positive relationship indicated that greater synergistic processing emerges from integration of correlated activity at synaptic timescales.

**Figure 2.**
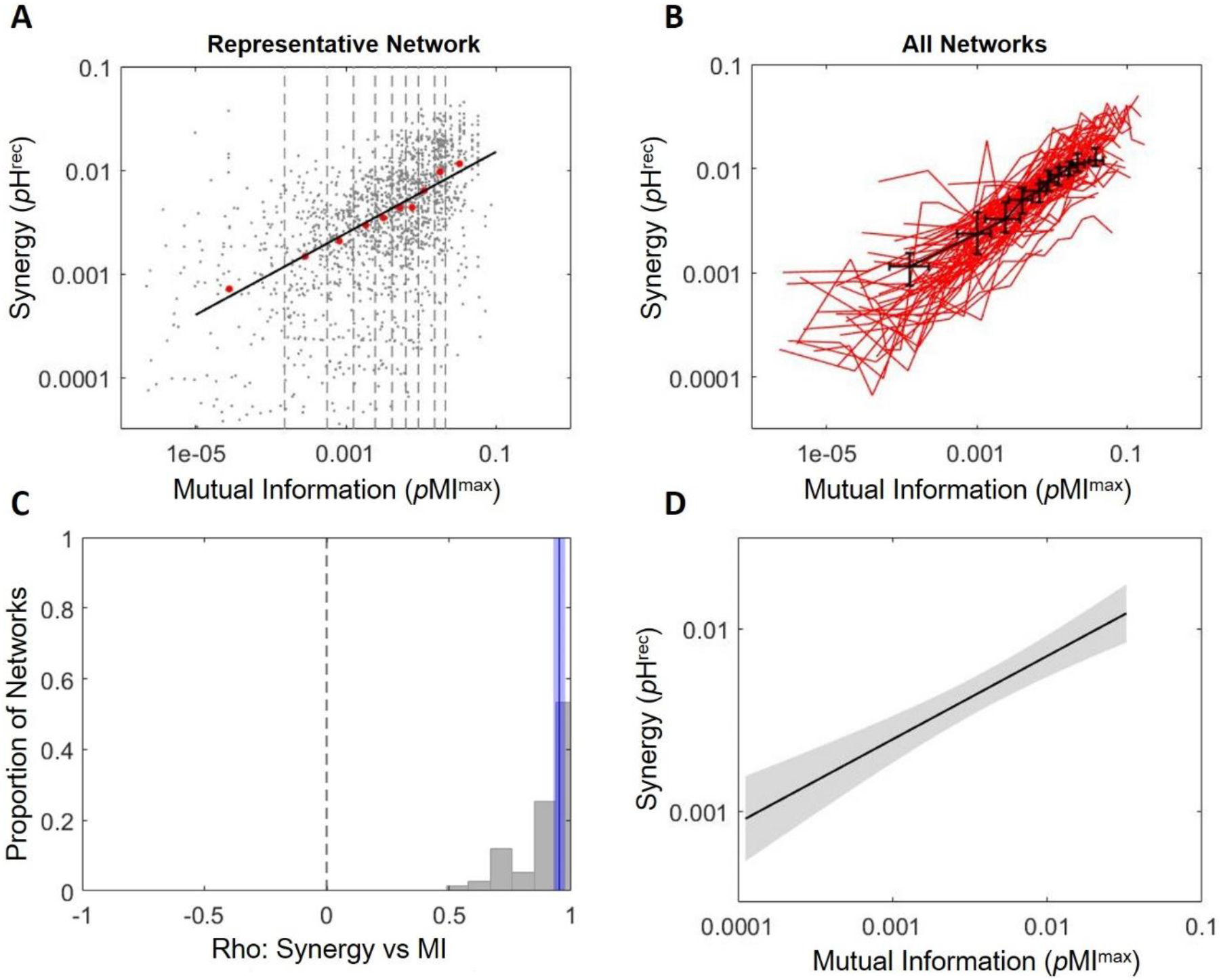
Increased correlated activity predicts greater synergistic processing at synaptic timescales. (A) Synergy plotted as a function of mutual information for a representative network shows a positive relationship. Grey dots indicate individual triads. Boundaries of deciles of mutual information are depicted via dashed lines. Median synergy for each decile (red dots) increased across deciles. The black line shows the linear least squares fit of the red dots (rho = 0.99, slope = 0.39). (B) Synergy across mutual information deciles for all 75 synaptic timescale networks are shown as red lines. The median synergy and mutual information for each decile is shown by the black points with error bars indicating 95% bootstrap confidence intervals. (C) Histogram of Spearman rank correlation coefficients for synergy versus mutual information shown in B. A solid blue line marks the median correlation coefficient across the 75 networks and the shaded region around the blue line shows the 95% bootstrap confidence interval around the median. (D) Average regression fit line across the 75 linear regressions of synergy versus mutual information indicates consistently positive relationship. Shaded region indicates the 95% confidence interval.

### Correlated activity is associated with greater gains in information transmission than redundancy at synaptic timescales

Theories regarding the favorability of correlated activity, or lack thereof, for neural computation are motivated by the impact of increasing correlation on information transmission and redundancy. To address these theories, we examined how information transmission and redundancy covaried with mutual information and how these relationships related to the generation of synergy. Information transmission was quantified as the multivariate transfer entropy (mvTE). We normalized mvTE to reflect the proportion of the entropy of the receiving neuron accounted for (*p*H^rec^). Redundancy was quantified using the redundancy term derived from partial information decomposition and was also normalized to reflect the proportion of the entropy of the receiving neuron accounted for (*p*H^rec^).

Theories that view correlated activity as favorable predict that correlated activity increases information transmission. Consistent with this, we found that mvTE was reliably positively correlated with mutual information at synaptic timescales (Spearman *r* = 0.99 [0.98 1.00]; Z_s.r._ = 7.58, n=75 networks, p <1×10^−13^; slope = 0.35 [0.32 0.38] Δlog_10_(*p*H^rec^) / Δlog_10_(*p*MI^max^); Fig 3A; Table 1). Theories that view correlated activity as unfavorable predict that correlated activity increases redundancy. Supporting this, we found that redundancy was significantly positively related to mutual information at synaptic timescales (Spearman *r* = 0.99 [0.98 0.99], Z_s.r._ = 7.56, n=75 networks, p <1×10^−13^; slope = 0.51 [0.47 0.57] Δlog_10_(*p*H^rec^) / Δlog_10_(*p*MI^max^); Fig 3A; Table 1). These results indicate that both hypotheses are well motivated with respect to the expected effect of increasing correlated activity.

**Table 1.** Relationships between information terms at synaptic timescales. Off diagonal: Median Spearman rank correlation coefficients (below diagonal) and linear regression slopes (above diagonal) for pairwise comparisons of synergy, redundancy, multivariate transfer entropy (mvTE), and the difference between mvTE and redundancy (mvTE-Red). Slopes are in units of Δ log_10_(*p*H^rec^) / Δ log_10_(*p*H^rec^). Along diagonal: Median Spearman rank correlation coefficients (left) and linear regression slopes (right) for each information term compared to mutual information. Slopes are in units of Δ log_10_(*p*H^rec^) / Δ log_10_(*p*MI^max^). * Indicate values that are significantly different from zero after Bonferroni-Holm correction (all *p* values < 0.001).

| Rho \ Slope | Synergy | Redundancy | mvTE | mvTE-Red |
| --- | --- | --- | --- | --- |
| Synergy | 0.95* \ 0.46* | 0.90* | 1.20* | 1.26* |
| Redundancy | 0.99* | 0.99* \ 0.51* | 1.34* | 1.41* |
| mvTE | 0.95* | 0.98* | 0.99* \ 0.35* | 1.09* |
| mvTE-Red | 0.89* | 0.93* | 0.99* | 0.96* \ 0.31* |

**Figure 3.**
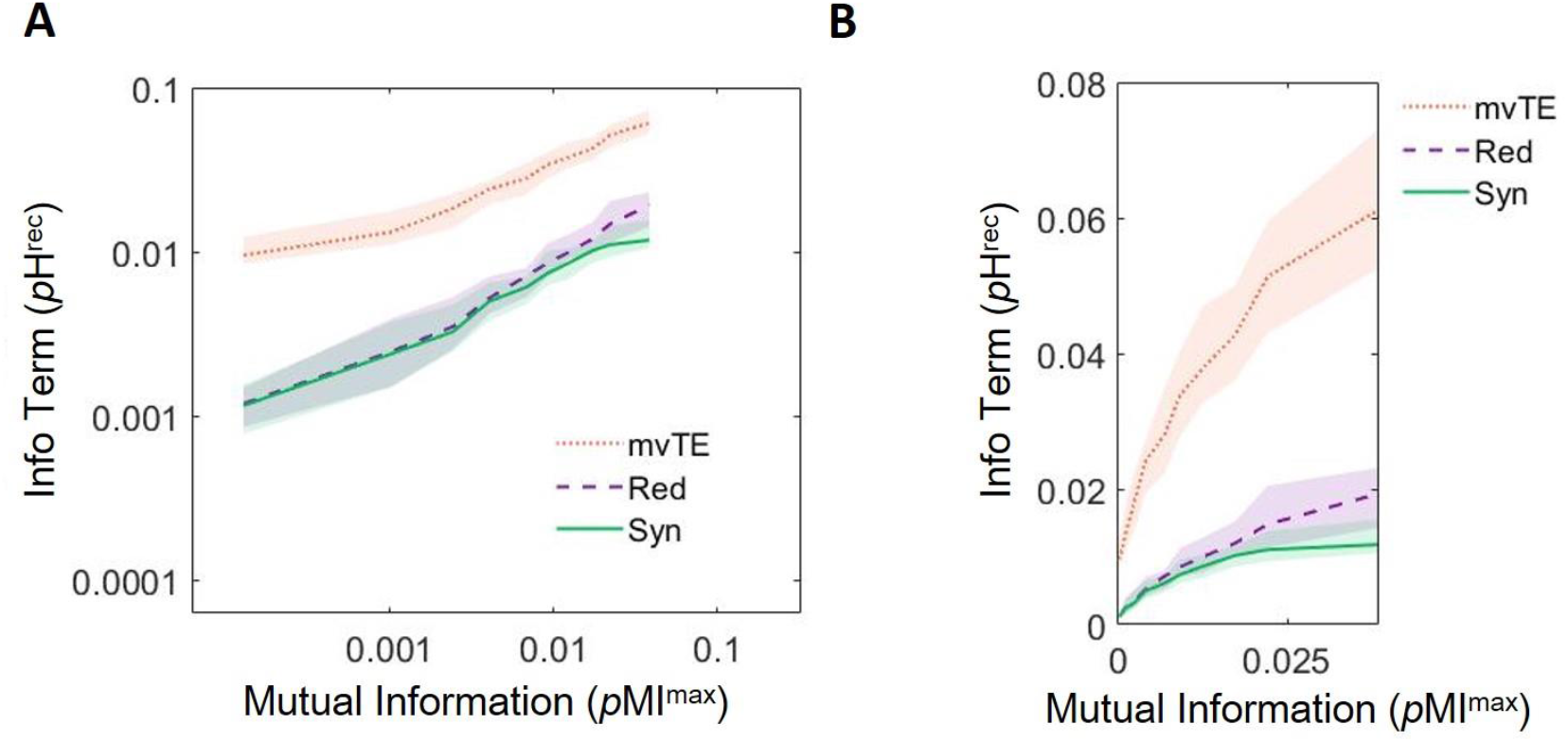
Growth of information transmission (mvTE) is greater than growth of redundancy over deciles of mutual information at synaptic timescales. (A) Multivariate transfer entropy (mvTE) and redundancy (Red) increase with mutual information. Synergy data is replotted here from Figure 2B for ease of comparison. (B) Same data as shown in A without log-scaling to show rapid growth of mvTE relative to redundancy. For all plots, lines depict medians and shaded regions depict 95% bootstrap confidence intervals around the median.

Next, we related the changes in mvTE and redundancy observed over deciles of Mutual information to those observed in synergy. Consistent with the idea that correlated activity can support greater computation by facilitating information transmission, synergy and mvTE were reliably positively correlated across deciles of mutual information (Spearman *r* = 0.95 [0.92 0.98]; Z_s.r._ = 7.53, p <1×10^−13^, n=75 networks; Table 1). Interestingly, synergy and redundancy were also reliably positively correlated across deciles of mutual information (Spearman *r* = 0.99 [0.98 0.99]; Z_s.r._ = 7.56, p <1×10^−13^, n = 75 networks; Table 1). This positive correlation is contrary to the idea that redundancy impedes computation at synaptic timescales.

We then examined the *relative* influence of correlated activity on information transmission and redundancy. In a non-log-scaled space, the slope relating mutual information to mvTE was significantly greater than the slope relating mutual information to redundancy (1.41 [1.26 1.58] vs. 0.48 [0.45 0.55] (Δ *p*H^rec^ / Δ *p*MI^max^); Z_s.r._ = 7.52, n=75 networks, p <1×10^−13^; Fig 3B). Note, though the slope relating Mutual information to mvTE was less than the slope relating mutual information to redundancy in the log-log space, mvTE was offset vertically over redundancy, corresponding to a significantly larger y-intercept (0.017 vs. 0.003, Z_s.r._ = 7.52, n=75 networks, p < 1×10^−11^; Fig 3A). This means that the exponential growth of mvTE was steeper than that of redundancy over the values of mutual information observed at synaptic timescales. This can be seen by comparing the relationship between mutual information and mvTE and redundancy without log-scaling (Fig 3B). Because mvTE increases faster than redundancy as a function of mutual information, the difference between the two also grew as a function of mutual information (slope = 0.13 [0.06 0.19] Δlog_10_(*p*H^rec^) / Δlog_10_(*p*MI^max^); Z_s.r._ = 4.19, p <1×10^−4^, n=75 networks; Table 1; curve not shown). The growth of this difference could account for how correlated activity supports synergistic processing at synaptic timescales, as it was positively correlated with synergy across deciles of mutual information (Spearman *r* = 0.88 [0.85 0.90]; Z_s.r._ = 7.52, p <1×10^−13^, n = 75 networks; Table 1).

### Activity becomes more correlated at extrasynaptic timescales, with negative returns for synergy

At synaptic timescales, (normalized) mutual information was generally low (~0.8% of maximum possible, as described above). From this, we reasoned that the observed positive relationship between mutual information and synergistic processing may have been due to synaptic processing operating in a regime of low overall correlation. To explore this further, we sought to test what happens in a regime of greater overall correlation.

We had observed that the median mutual information increased progressively across the three synaptic timescales (*p*MI^max^ across timescales: 3 ms = 0.004 [0.002 0.005]; 5 ms = 0.008 [0.006 0.010]; & 11 ms = 0.019 [0.013 0.023]; Figure 4A), suggesting that extending the analysis to longer timescales would yield even greater mutual information. Examining median mutual information in seven additional logarithmically spaced timescales, extending out to 3000 ms, showed that it indeed continued to increase (*p*MI^max^ across timescales: 23 ms = 0.03 [0.02 0.04]; 48 ms = 0.07 [0.05 0.08]; 104 ms = 0.12 [0.09 0.17]; 225 ms = 0.16 [0.10 0.23]; 485 ms = 0.20 [0.14 0.26]; 1044 ms = 0.22 [0.17 0.27]; 2250 ms = 0.25 [0.21 0.33]). All data is shown in Fig 4B.

**Figure 4.**
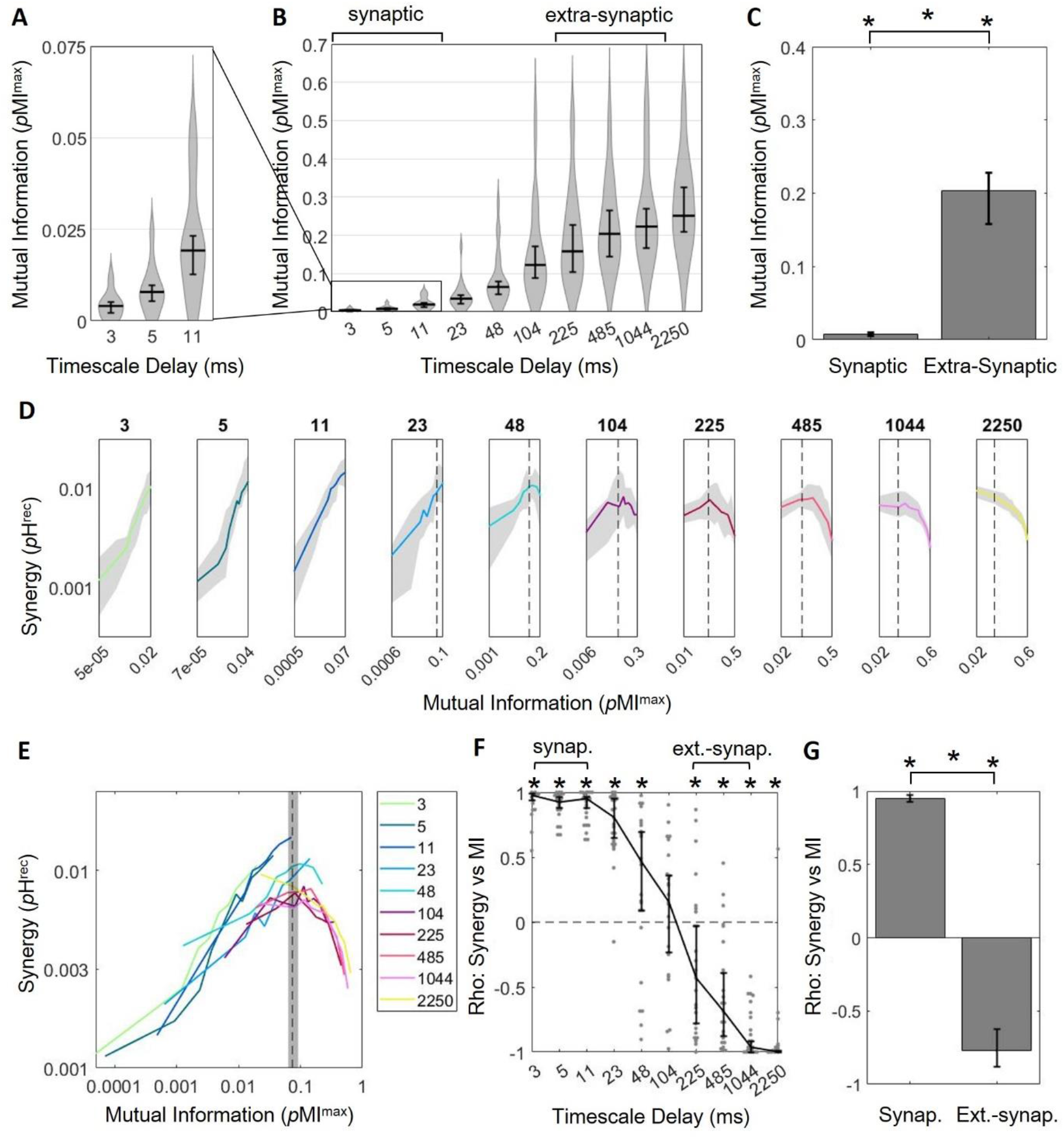
Greater correlated activity emerges at longer timescales with diminishing returns for synergistic processing. (A) Median mutual information increases across synaptic timescales. (B) Median mutual information continues to increase at extrasynaptic timescales. (C) Median mutual information is significantly larger at extrasynaptic timescales than at synaptic timescales. (D) Plotting synergy as a function of mutual information across timescales shows that the positive relationship between synergy and mutual information only exists at synaptic timescales where mutual information is relatively low. This relationship becomes negative at extrasynaptic timescales where mutual information is high. Note that mutual information, plotted along the x-axis, varies across sub-panels. A dashed vertical line where mutual information is 0.07 is included to facilitate visual alignment across panels. (E) Curves from D are replotted on the same axes here to show the overall relationship between synergy and mutual information. (F) Spearman rank correlation coefficients of synergy versus mutual information for all networks across timescales show that synergy and mutual information are positively related at shorter timescales and negatively related at longer timescales. (G) Synergy and mutual information are significantly positively correlated at synaptic timescales and significantly negatively correlated at extra-synaptic timescales. For all plots shown above, bars/solid lines indicate medians, shaded regions/error bars depict 95% bootstrap confidence intervals around the median, and * indicates p < 0.05.

To enable statistical comparison between the patterns observed at longer timescales with those we reported for the synaptic timescale, we defined the timescales that were two orders of magnitude longer than those used to span the synaptic timescales as the ‘extrasynaptic timescale’ (i.e., those centered on 225, 485, & 1044 ms versus 3, 5, & 11 ms). Comparing mutual information between these two timescales revealed that correlated activity was significantly greater at extrasynaptic timescales than synaptic timescales (0.20 [0.16 0.23] vs. 0.008 [0.005 0.010] *p*MI^max^, Z_r.s._ = 10.51, n = 150 networks, p <1×10^−25^; Fig 4C).

Analyzing the relationship between mutual information and synergy at longer timescales revealed that the positive correlation observed at synaptic timescales was lost as the median mutual information increased further (Fig 4D-G). At timescales beyond the synaptic timescales, the relationship between mutual information and synergy inverted gradually. Numerically, this can be seen in the Spearman rank correlation coefficients relating mutual information to synergy, which started positive and gradually shifted to negative across timescales (Fig 4F; Spearman *r* across timescales: 3 ms = 0.98 [0.94 0.99]**; 5 ms = 0.93 [0.88 0.96]**; 11 ms = 0.95 [0.88 0.96]**; 23 ms = 0.81 [0.65 0.95]**; 48 ms = 0.47 [0.09 0.70]**; 104 ms = 0.15 [−0.24 0.36]; 225 ms = −0.43 [−0.78 −0.03]**; 485 ms = −0.68 [−0.88 −0.39]**; 1044 ms = −0.96 [−1.00 −0.92]**; 2250 ms = −1.00 [−1.00 −0.99]**; ** = p <0.05). Though the overall trend between mutual information and synergy remained positive at the ~48 ms timescale, evidence of a saturation point emerged as synergy began to decrease for mutual information values greater than ~ 0.07. Across networks, the saturation point was reliably around 0.07 [0.06 0.09]. At longer timescales, a greater percentile of triads had mutual information values above 0.07. This resulted in diminishing returns of synergy as mutual information grew further (Fig 4D-E). The relationship between mutual information and synergy was significantly negative at timescales including and greater than 225 ms. Whereas mutual information and synergy were positively correlated at synaptic timescales (as described above), mutual information was significantly negatively correlated with synergy at extrasynaptic timescales (Spearman *r* = −0.77 [−0.88, −0.62]; Z_s.r._ = −6.57, n = 75 networks, p <1×10^−10^; slope= −0.16 [−0.21 −0.10] Δlog_10_(*p*H^rec^) /Δlog_10_(*p*MI^max^); Fig 4F-G). The slopes observed at the extrasynaptic timescales were significantly less than those observed at synaptic timescales (Zr.s, = −10.40, n = 150 networks, p <1×10^−24^).

These results were obtained using mutual information and synergy values that were normalized by the maximum possible values given the firing rates and bin sizes for each triad at each timescale. This was done because information terms such as mutual information and synergy are sensitive to the entropy of the data being analyzed, which is, in turn, sensitive to firing rates and bin sizes. The log-normal distribution of firing rates and the different timescales generated variability in the total entropy and thus information terms. By normalizing, we minimized any effects that may have been due to changes in entropy within and across timescales. Critically, however, none of the qualitative results described here depended upon this normalization (see Supplemental Figure 6 for non-normalized results).

### Correlated activity is associated with greater gains in redundancy than information transmission at extrasynaptic timescales

The robust positive relationship between mvTE and mutual information observed at synaptic timescales were lost at longer timescales (Fig 5A-B). This can be seen in the Spearman rank correlation coefficients relating mvTE to mutual information, which started positive and gradually shifted towards zero across timescales (Fig 5C; Spearman *r* across timescales: 3 ms = 0.99 [0.96 1.00]**; 5 ms = 0.98 [0.93 1.00]**; 11 ms = 0.99 [0.96 1.00]**; 23 ms = 0.98 [0.93 1.00]**; 48 ms = 0.90 [0.28 0.98]**; 104 ms = 0.95 [0.89 0.99]**; 225 ms = 0.99 [0.94 1.00]**; 485 ms = 0.88 [0.42 0.95]**; 1044 ms = 0.27 [−0.39 0.71]; 2250 ms = −0.25 [−0.72 0.54]; ** indicates p <0.05). Consistent with the loss of a relationship between mvTE and mutual information at longer timescales, mvTE ceased to be reliably correlated with synergy across deciles of mutual information at extrasynaptic timescales (Spearman *r* = −0.19 [−0.41 0.03]; Z_s.r._ = −1.88, p = 0.06, n=75 networks; Fig 5D). The correlation between mvTE and synergy was significantly greater at synaptic compared to extra-synaptic timescales (Z_r.s._ = 9.99, p <1×10^−22^, n = 150 networks; Fig 5D). These results indicate that the growth of information transmission with correlated activity that contributed to increased synergistic processing at synaptic timescales was lost at extrasynaptic timescales.

**Figure 5.**
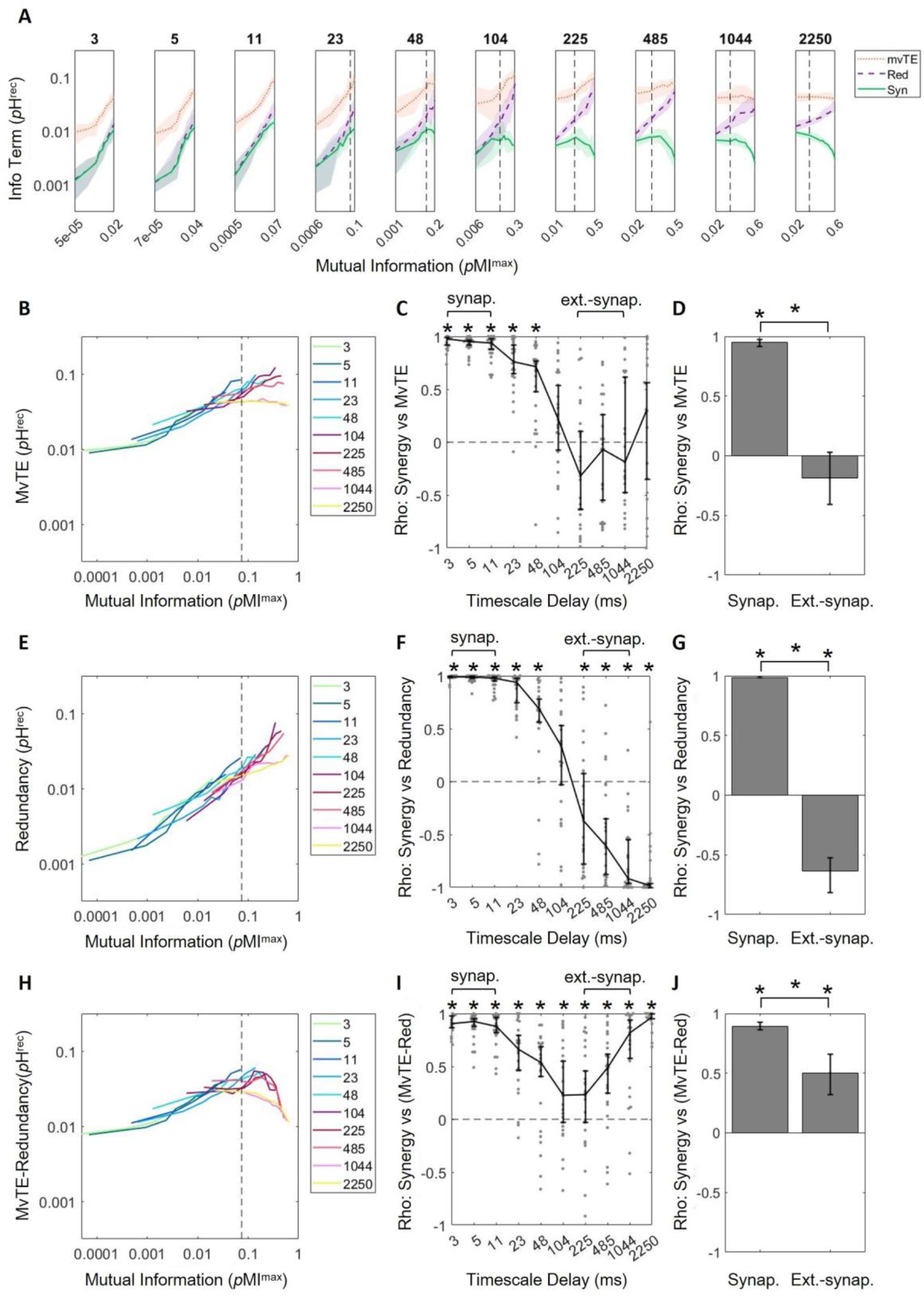
Redundancy encroaches upon multivariate transfer entropy at longer timescales, accounting for decreasing synergy at those timescales. (A) Log-scaled multivariate transfer entropy (mvTE), Redundancy (Red), and Synergy (Syn) versus mutual information, for all timescales. Growth of mvTE with respect to mutual information is steep at short timescales, but decreases across timescales. Redundancy grows with mutual information at all timescales. (B) MvTE curves in A plotted on the same axes. (C) Spearman rank correlation coefficients of synergy versus mvTE across timescales indicates that mvTE saturates at longer timescales. (D) Synergy and mvTE are significantly positively related at synaptic timescales, but not at extra-synaptic timescales. (E) Redundancy curves in A plotted on the same axes. (F) Spearman rank correlation coefficients of synergy versus redundancy across timescales shows how the two are positively related at shorter timescales and negatively related at longer timescales. (G) Synergy and redundancy are significantly positively related at synaptic timescales and significantly negatively related at extra-synaptic timescales. (H) Difference of MvTE and Red curves in A plotted on the same axes shows that the overall trend looks most similar to that of synergy versus MI in Figure 4E. (I) Spearman rank correlation coefficients of synergy versus mvTE-redundancy across timescales shows that the two are positively related at all timescales. (J) Synergy and mvTE-redundancy are significantly positively related at synaptic and extrasynaptic timescales. Synergy versus mutual information data has been replotted from Fig 4D here in A for ease of comparison. For all plots, solid/bold lines indicate medians and shaded regions/error bars depict 95% bootstrap confidence intervals around the median.

Redundancy remained significantly positively related with mutual information at all timescales (Fig 5E; Spearman *r* across timescales: 3 ms = 0.98 [0.94 0.99]**; 5 ms = 0.98 [0.92 0.99]**; 11ms = 0.99 [0.95 1.00]**; 23 ms = 0.99 [0.96 1.00]**; 48 ms = 0.95 [0.87 0.99]**; 104 ms = 0.99 [0.96 1.00]**; 225 ms = 1.00 [0.99 1.00]**; 485 ms = 1.00 [0.99 1.00]**; 1044 ms = 1.00 [0.98 1.00]**; 2250 ms = 1.00 [1.00 1.00]**; ** indicates p <0.05). Though synergy and redundancy had been positively correlated at synaptic timescales (as described above), we found that they were negatively correlated at extrasynaptic timescales (Spearman *r* = −0.64 [−0.82 −0.53]; Z_s.r._ = −5.95, p <1×10^−8^; n=75 networks; Fig 5G). The correlation between redundancy and synergy was significantly greater at synaptic compared to extra-synaptic timescales (Z_r.s._ = 10.55, p <1×10^−25^, n = 150 networks; Fig 5G). This result indicates that, at extrasynaptic timescales, redundancy impeded synergistic processing.

Because redundancy, but not mvTE, continued to increase as a function of mutual information at longer timescales, the difference between the two became smaller. This difference continued to be positively correlated with synergy at extrasynaptic timescales (Spearman *r* = 0.50 [0.32 0.66]; Z_s.r._ = 5.60, p <1×10^−7^; n=75 networks; Fig 5J). The correlation between mvTE-redundancy and synergy was significantly greater at synaptic compared to extra-synaptic timescales (Z_r.s._ = 6.57, p <1×10^−10^, n = 150 networks; Fig 5J), however, correlations were significantly positive at both timescales. This indicated that reduction of the gap between mvTE and redundancy across timescales could account for why correlated activity stops supporting synergistic processing at extrasynaptic timescales.

## DISCUSSION

Our aim with the work described here was to determine whether correlated activity in cortical microcircuits is favorable or unfavorable for synergistic processing. By applying information theoretic analyses to the spiking activity of hundreds of neurons, we showed that correlated activity is favorable for synergistic processing when the overall level of correlations are low. We also showed that correlated activity is unfavorable for synergistic processing when overall correlation levels were an order of magnitude greater. Secondary analyses indicated that the relationship between correlated activity and synergistic processing varied as a function of the relative amount of information transmission and redundancy. When overall correlation levels were low, we observed that incremental increases in correlation were associated with larger increases in information transmission than redundancy. We hypothesize that this effectively left greater bandwidth available for synergistic processing. When overall correlation levels were high, incremental increases in correlation were associated with larger increases in redundancy than information transmission. We hypothesize that this effectively caused redundancy to consume the bandwidth available for synergistic processing. This is summarized in Figure 6.

**Figure 6.**
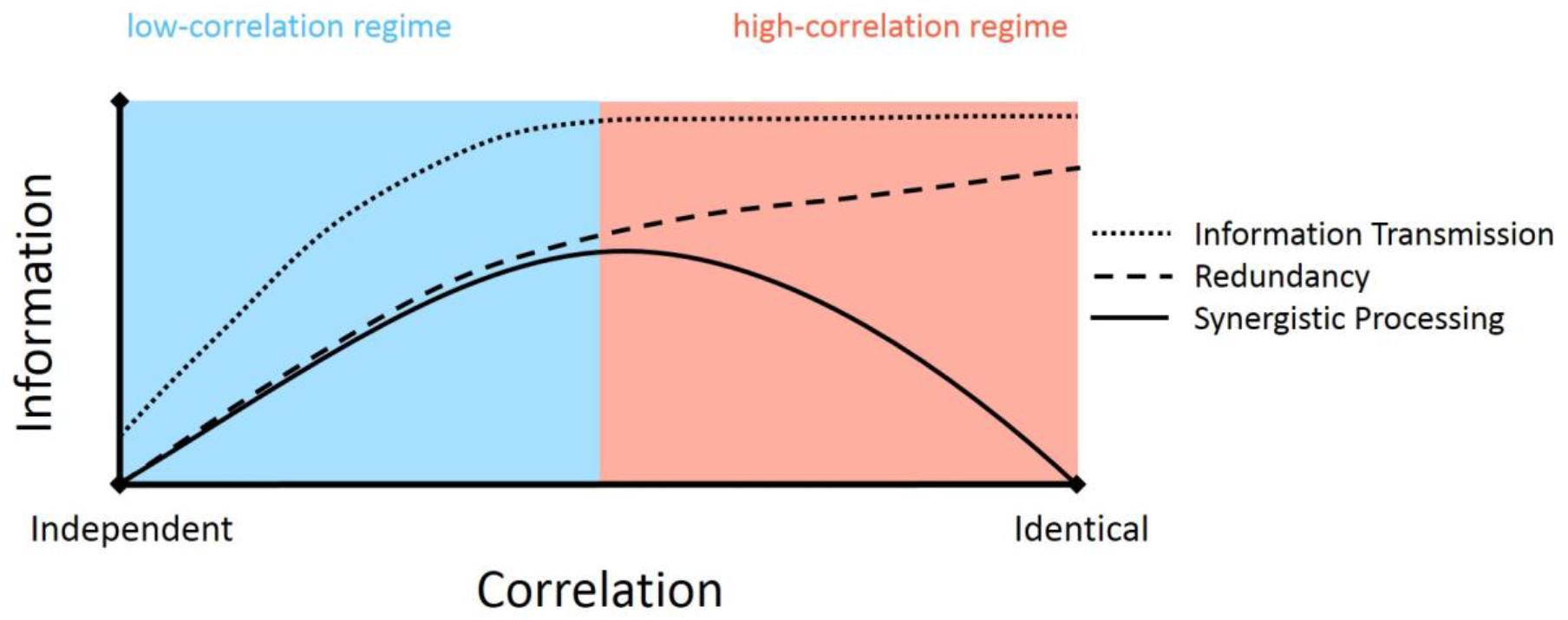
Neural computation can be expressed as a function of the correlation between inputs. When the correlation of inputs is low, synergistic processing benefits from increased correlation. In this regime, both information transmission and redundancy increase with greater correlation but information transmission grows faster than redundancy effectively increasing the bandwidth available for synergistic processing. When the correlation of inputs is high, synergistic processing is hurt by increased correlation. In this regime, redundancy and not information transmission increases with greater correlation, reducing the bandwidth available for synergistic processing.

### Correlation and information processing emerge with cognition

Correlated activity has been shown to increase as cognition emerges. This has been shown, both during neuronal development (Chiappalone et al., 2006) and during the transition out of deep anesthesia (Bettinardi et al., 2015). Recovery from anesthesia also brings about an increase in information transmission between neurons (Fagerholm et al., 2016). Given the positive relationship we describe here, we predict that computation would also increase under these contexts. Further, we predict that it is the increase in computation that marks the emergence of cognition. We propose this because without computation, or information modification events in which the whole is greater than the sum of the parts, all neuronal activity would be passive information transfer, which cannot account for the emergence of higher order brain states and behavior. To establish this more concretely, future studies should examine synergy in the context of cognition and criticality.

### Correlation regimes parse the favorability debate

These findings provide empirical support for each of two seemingly contradictory perspectives regarding the favorability of correlated activity for neural information processing. The first perspective, held by many in the cognitive rhythms community, states that correlated activity is favorable for information processing because correlated activity facilitates information transmission (Averbeck & Lee, 2004; Fries, 2005; Fries, 2015; Lisman & Jensen, 2013; Salinas & Sejnowski, 2001). The second perspective, held by many in the sensory processing and artificial neural networks communities, states that correlated activity is unfavorable for information processing because correlated activity is viewed as equivalent to capacity-consuming redundancy (Attneave 1954; Barlow 1961). Even within the respective communities there is recognition that the relationship between correlated activity and processing may not be so straight-forward. For example, within the cognitive rhythms community, Hanslmayr has explicitly argued that synchronized activity can both facilitate and impede processing (Hanslmayr et al., 2012; Hanslmayr, Staresina, & Bowman, 2016). Likewise, in the sensory processing community, there is evidence that correlated activity can also increase processing (e.g., Nigam, Pojoga, & Dragoi, 2019). Each perspective, our data suggest, is relevant in distinct correlation regimes observed, here, across timescales.

The idea that the favorability of correlated activity for information processing varies non-monotonically can account for seeming inconsistencies both within and between the separate perspectives. For example, while correlated activity is largely held as benefiting cognitive processing within the cognitive rhythms community (e.g., Fries, 2005; Lisman & Jensen, 2013), it can be that information processing requires decorrelation in higher correlation regimes (e.g., Hanslmayr et al., 2012). Indeed, it has been suggested that correlated activity serves distinct roles in different neural systems such as the cortex and hippocampus (Hanslmayr, et al., 2016). Consistent with the possibility for regional differences in the favorability of correlated activity, the present findings indicate that the key differentiating functional factor between areas is the correlation level. Breaking from simple interregional differences, however, this framework would also predict that there can be systematic variability in the favorability of correlated activity even within a region. This is consistent with the finding that correlated activity improves synergistic coding in primary visual cortex (e.g., Nigam et al., 2019). As such, the present findings also serve to account for apparent inconsistencies in the sensory processing community.

### Correlation regime as a covariate of timescale

Importantly, the variance in correlation regime we observed was confounded with timescale. That is, we observed the low correlation regime at short timescales and the high correlation regime at long timescales. While our motivating question was grounded in the importance of correlation rather than timescale, it is not possible to disentangle these two factors fully here and additional work is needed on this topic.

### ‘Signal’ versus ‘noise’ correlations

The analysis of correlated activity used here differs from the approach commonly used in sensory processing and artificial neural networks research in that we did not separately analyze ‘signal correlations’ and ‘noise correlations.’ In such research, in which neural responses to experimentally delivered stimuli are evaluated, it is possible to separate activity into what is stimulus-dependent (aka, signal correlations) and what is internally generated (aka, noise correlations) (Cohen & Kohn, 2011; Gawne & Richmond, 1993). Here, no stimuli were delivered and the analysis integrated activity over the entire hour of the recording, likening our analysis to one of noise correlations only. This does not mean, however, that the non-monotonic function relating correlated activity to synergistic processing need apply only to noise correlations. Though signal and noise correlations can be analytically separated, the same underlying physiological constraints should be expected to hold for both types of correlations. If there is an important difference between the activity comprising signal-and noise-correlations, it is, we argue, that correlations induced by extrinsic factors (e.g., delivered stimuli) reflect transient shifts in the statistics of the local processing dynamics. To determine how these results relate to signal and noise correlations, future work must test how synergistic processing relates to the level of correlated activity in response to stimuli delivered *in vivo*.

### Related works

The present findings are consistent with prior analyses of synergistic processing and, we hypothesize, could account for previously reported topological determinants of synergistic processing. In a previous analysis of synergistic processing in developing organotypic cultures, Wibral et al. (2017) observed a similar pattern of synergy increasing initially, and then decreasing, as a function of culture maturity. Notably, they also found that mutual information among the neurons increased steadily across the same developmental timeline. Thus, the underlying pattern of results is consistent with the pattern observed here. Previous analysis of the topological determinants of synergistic processing found that synergistic processing increased with the number of connections made by upstream neurons (Timme et al., 2016). In line with this observation, we have previously shown that neurons in rich clubs of cortical micro-circuits do about twice as much synergistic processing as neurons outside of rich clubs (Faber et al., 2019). Based on our present findings, we predict that (1) upstream neurons with more connections have greater correlated activity, and (2) rich club neurons have greater correlated activity than neurons outside of the rich club.

### Significance of optimal synergistic processing

The non-monotonic curve relating mutual information to synergy revealed by our analyses indicates that the most synergistic processing was observed when the activity of the sending neurons was correlated at ~7% of the maximum possible. It is unclear whether this reflects a universal optimum or if this value should be expected to vary as a function of the local circuit physiology. Physiology plays an important role in regulating the amount of correlation (e.g., Gutnisky and Dragoi, 2008; Minces et al., 2017). What is not clear based on existing work, however, is whether these changes shift processing relative to a fixed local optimum or whether the optimal correlation also shifts. To address this, future research must compare how the optimal value varies across brain regions, physiological conditions, and behavioral conditions, if at all.

### Relevance of in vitro preparation

The use of organotypic cultures in the present work facilitated the recording of hundreds of neurons simultaneously. While organotypic cultures naturally differ from intact *in vivo* tissue, organotypic cultures nonetheless exhibit synaptic structure and electrophysiological activity very similar to that found *in vivo* (Beggs & Plenz, 2004; Bolz et al., 1990; Caeser et al., 1989; Götz & Bolz, 1992; Ikegaya et al., 2004; Klostermann &Wahle, 1999; Plenz & Aertsen, 1996). For example, the distribution of firing rates observed in cultures is lognormal, as seen *in vivo* (Nigam et al., 2016), and the strengths of functional connections are lognormally distributed, similar to the distribution of synaptic strengths observed in patch clamp recordings (reviewed in Buzsáki & Mizuseki, 2014; Song et al., 2005). These features indicate that organotypic cortical cultures serve as a reasonable model system for exploring local cortical networks, while offering unique accessibility to large neuron count and high temporal resolution recordings. However, additional work will need to be done to understand how the synergy-mutual information relationship observed *in vitro* differs from what may exist *in vivo*, particularly in the context of behavior.

While stimulus-driven activity has been favored in research for its ability to provide insight into neural coding mechanisms, such studies assume that the brain is primarily reflexive and that internal dynamics are not informative with regard to information processing. However, internally-driven *spontaneous* activity of neurons, or activity that does not track external variables in observable ways, has been repeatedly shown to be no less cognitively interesting than stimulus-linked activity (Johnson et al., 2009; Raichle, 2010; for a review see Tozzi et al., 2016; Tsodyks et al., 1999). Not only is spontaneous activity predominant throughout the brain, but it also drives critical processes such as neuronal development (Cang et al., 2005; Chiappalone et al., 2006; Wibral et al., 2017).

### Conclusions

We show, for the first time, that the relationship between correlated activity and synergistic processing is positive at synaptic timescales with low levels of overall correlation but negative at extrasynaptic timescales with high levels of overall correlation. We propose that the relationship between correlated activity and synergistic processing varies as a function of the overall correlation regime. The shift from a positive to a negative relationship between regimes in this framework is accounted for by shifts in the effects of increasing correlation on information transmission and redundancy. In the low correlation regime, increases in correlation increased information transmission faster than redundancy, thereby increasing the effective bandwidth for synergistic processing. In the high correlation regime, increases in correlation increased redundancy faster than information transmission, thereby decreasing the effective bandwidth for synergistic processing. Both dynamics had been hypothesized to exist previously by separate frameworks within separate research communities. The present work serves to bridge across these disconnected domains of work to synthesize the two perspectives into a unified framework.

## MATERIALS & METHODS

To answer the question of how correlation and computation are related in cortical circuits, we combined network analysis with information theoretic tools to analyze the spiking activity of hundreds of neurons recorded from organotypic cultures of mouse somatosensory cortex. Due to space limitations, here we provide an overview of our methods and focus on those steps that are most relevant for interpreting our results. A comprehensive description of all our methods can be found in the Supplemental Materials.

All procedures were performed in strict accordance with guidelines from the National Institutes of Health, and approved by the Animal Care and Use Committees of Indiana University and the University of California, Santa Cruz.

### Electrophysiological recordings

All results reported here were derived from the analysis of electrophysiological recordings of 25 organotypic cultures prepared from slices of mouse somatosensory cortex. One hour long recordings were performed at 20 kHz sampling using a 512-channel array of 5 μm diameter electrodes arranged in a triangular lattice with an inter-electrode distance of 60 μm (spanning approximately 0.9 mm by 1.9 mm). Once the data were collected, spikes were sorted using a PCA approach (Ito et al., 2014; Litke et al., 2004; Timme et al., 2014) to form spike trains of between 98 and 594 (median = 310) well isolated individual neurons depending on the recording.

### Network construction

Networks of effective connectivity, representing global activity in recordings, were constructed following the methods described by Timme et al. (2014, 2016). Briefly, weighted effective connections between neurons were established using transfer entropy (TE; Schreiber, 2000). To consider synaptic interactions, we computed TE at three timescales spanning 0.05 – 14 ms, discretized into overlapping bins of 0.05-3 ms, 1.6-6.4 ms, and 3.5-14 ms. To consider extrasynaptic interactions, we computed additional networks at timescales spanning 7.5-3000 ms, discretized into overlapping bins of 7.5-30 ms, 16.15-64.6 ms, 34.8-139.2 ms, 75-300 ms, 161.6-646.4 ms, 348.1-1392.4 ms, and 750-3000 ms (Supplemental Table 1). This resulted in an additional 7 networks per recording, or 175 networks (250 networks total). Only significant TE, determined through comparison to the TE values obtained with jittered spike trains (α = 0.001; 5000 jitters), were used in the construction of the networks. TE values were normalized by the total entropy of the receiving neuron so as to reflect the proportion of the receiver neuron’s capacity that can be accounted for by the transmitting neuron.

### Quantifying correlation

The degree of similarity between all pairs of neurons was quantified as mutual information (Cover & Thomas, 1991; Shannon & Weaver, 1949):

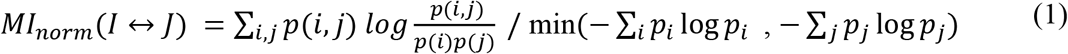

Mutual information is a symmetric measure which quantifies how much uncertainty is reduced in one variable (I) by knowing another (J). All mutual information values were normalized by the maximum possible mutual information that could have occurred between each pair of neurons, which is equivalent to the minimum entropy of the two neurons (Gray & Shields, 1977). Using the normalized values allowed us to consider the proportion of the total possible mutual information that occurred between neurons and to control for differences in entropy across pairs of neurons, improving the interpretability of the numbers obtained. The results reported here are robust to the choice of similarity measure, as shown in Supplemental Figures 4 and 5.

### Quantifying computation, redundancy, and information transmission

Computation was operationalized as synergy. Synergy measures the additional information regarding the future state of the receiver, gained by considering the prior state of the senders jointly, beyond what they offered individually, after accounting for the redundancy between the sending neurons and the past state of the receiver itself. Synergy was calculated according to the partial information decomposition (PID) approach described by Williams and Beer (2011), including use of the *I*_min_ term to calculate redundancy (see Supplemental Material). PID compares the measured bivariate TE between neurons *TE*(*J*→*I*) and *TE*(*K*→*I*) with the measured multivariate TE (the triad-level information transmission) among neurons *TE*({*J,K*}→*I*) to estimate terms that reflect the unique information carried by each neuron, the redundancy between neurons, and the synergy (i.e., gain over the sum of the parts) between neurons. Redundancy was computed as per Supplemental equations 8-10. Synergy was then computed via:

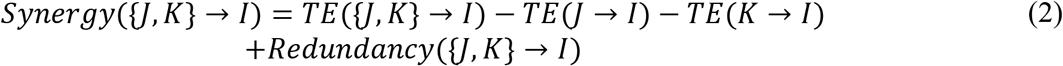

As with bivariate TE, synergy, redundancy and multivariate TE (mvTE) were normalized by the total entropy of the receiving neuron.

Although there are other methods for calculating partial information terms (Bertschinger et al., 2014; Lizier et al., 2018; Pica et al., 2017; Wibral et al., 2017), we chose this measure because it is capable of detecting linear and nonlinear interactions and it has been shown to be effective for our datatype (Timme et al., 2016; Faber et al., 2019). In addition, unlike other methods (Lizier et al., 2011; Stramaglia et al., 2012), PID of mvTE can decompose the interaction into non-negative and non-overlapping terms. However, to address previously raised concerns that PID overestimates the redundancy term (Bertschinger et al., 2014; Pica et al., 2017), and consequently synergy, we also used an alternate implementation of PID that estimates synergy based on the lower bound of redundancy. In this implementation, the effective threshold for triads to generate synergy is higher. This approach yielded the same qualitative pattern of results at synaptic timescales (see Supplemental Figure 7). At extra-synaptic timescales, this alternate version of synergy remained positively related to mutual information. This is likely due to the fact that the increased threshold meant that only triads with larger amounts of synergy were included in this analysis, which also meant that redundancy had not effectively cannibalized the mvTE.

Note, we did not examine interactions larger than triads due to the multi-fold increase in the computational burden that arises in considering higher order synergy terms. In addition to the combinatorial explosion of increased numbers of inputs, the number of PID terms increases rapidly as the number of variables increases. However, based on bounds calculated for the highest order synergy term by Timme et al. (2016), it was determined that the information gained by including an additional input beyond two either remained constant or decreased. Thus, it was inferred that lower order (two-input) computations dominated. In addition, although we did not consider more than two inputs at a time, because we considered all possible triads in each network, we effectively sub-sampled the entire space of inputs for each neuron.

Also, note that mutual information (or any type of correlation) between senders does not necessarily imply that sender neurons have redundant information about the receiver neuron; although it does make redundancy more likely. Of course, for large mutual information values, redundancy becomes inevitable, as the majority of the information between the senders is shared, and therefore any information they may have about the receiver is likely to be overlapping.

### Statistics

To aggregate results across networks with differing numbers of triads and to maintain the ability to observe possible non-monotonic relationships between sender mutual information and synergy, we collapsed synergy, redundancy and mvTE across triads based on deciles of sender mutual information for each network. That is, we calculated the median values of synergy, redundancy, and mvTE for each decile of sender mutual information. These values were used in all analyses. We obtained the same qualitative pattern of results when analyses are performed at the triad-level.

All values were log_10_ scaled in all analyses unless stated otherwise. This scaling was motivated by the inherent log-normality of these variables, as well as by the goodness-of-fit obtained for relationships in the log_10_ space compared to the linear space.

All results are reported as medians followed by the 95% bootstrap confidence limits (computed using 10,000 iterations) reported inside of square brackets. Accordingly, figures depict the medians with errorbars reflecting the 95% bootstrap confidence limits. Comparisons between conditions or against null models were performed using the nonparametric Wilcoxon signed-rank test, unless specified otherwise. The threshold for significance was set at 0.05, unless indicated otherwise in the text. Bonferroni-Holm corrections were used in cases of multiple comparisons.

## ACKNOWLEDGEMENTS

We thank Benjamin Dann, Colleen Hughes, and Thomas Kreuz for helpful comments and discussion.

## FUNDING INFORMATION

Ehren L. Newman, Whitehall Foundation (http://dx.doi.org/10.13039/100001391), Award ID: 17-12-114. John M. Beggs, National Science Foundation (http://dx.doi.org/10.13039/100000001), Award ID: 1429500. John M. Beggs, National Science Foundation (http://dx.doi.org/10.13039/100000001), Award ID: 1513779. Samantha P. Faber, National Science Foundation (http://dx.doi.org/10.13039/100000001), Award ID: 1735095; Samantha P. Faber, Indiana Space Grant Consortium.

