## Supplemental Materials for "Correlated activity favors synergistic processing in local cortical networks *in vitro* at synaptically-relevant timescales"

SUPPLEMENTAL METHODS

The methods used in this study are described below in the following order: (1) in vitro data collection; (2) construction of effective connectivity networks; and (3) quantification of neural computation and related information measures.

***In vitro* data collection**

All procedures were performed in strict accordance with guidelines from the National Institutes of Health, and approved by the Animal Care and Use Committees of Indiana University and the University of California, Santa Cruz.

To study the relationship between neural computation and topological measures of networks of spiking neurons, we analyzed data collected *in vitro*. Data were spontaneously spiking organotypic cultures of mouse somatosensory cortex obtained from postnatal Day 6 to 7 Black 6 mouse pups (RRID:Charles_River:24101632, Harlan) following methods described by Tang et al. (2008) & Ito et al. (2014). Spontaneous (as opposed to stimulus-driven) spiking activity in the cultures was recorded at a high temporal resolution of 50 μs, between 2 and 4 weeks after culture preparation, using a 512-microelectrode array (Litke et al., 2004). Array electrodes were flat, 5 μm in diameter and arranged in a triangular lattice with an interelectrode distance of 60 μm. This spacing means that the spiking of most cells is picked up by multiple sites and there are few gaps where cells are too far from electrodes to be recorded. The full array allowed for a total recording area of approximately 0.9 mm by 1.9 mm. This preparation and recording method enabled the isolation of large numbers of neurons (a median of 310 cells per recording in 25 hour-long recordings) at high temporal resolution, beyond what can currently be done in any *in vivo* setup. Crucially, the temporal resolution of this method was small enough to resolve synaptic delays typically found in cortex (Mason et al., 1991; Swadlow, 1994).

Once the data were collected, spikes were sorted using a PCA approach based on waveforms detected at seven adjacent electrodes (Ito et al., 2014; Litke et al., 2004; Timme et al., 2014). This process yielded a single set of spike times for each isolated neuron. Neurons that spiked fewer than 100 spikes during the hour long recording were removed from the analysis. Spike trains were then used to build networks.

**Effective connectivity network construction**

Because neural computation is fundamentally a dynamic process, we focused on examining networks of effective connectivity. In these networks, connections represent a predictive relationship between the firing of two different neurons. Note, effective connectivity differs from structural connectivity (synapses or gap junctions between neurons) and functional connectivity (e.g., cross-correlations between neuronal time series). Here, effective connections represent directed information transfer between neurons.

Networks of effective connectivity, representing global activity in recordings, were constructed following methods described previously (Timme et al. 2014 & 2016) using a measure from information theory known as transfer entropy (TE; Schreiber, 2000). TE was selected for its ability to detect nonlinear interactions and deal with discrete data, such as spike trains. To capture neuron interactions at timescales relevant to synaptic transmission (1-14 ms; Mason et al., 1991; Swadlow, 1994), multiple windows are used to improve the sensitivity to functional interactions across these delays. Spiking data was binned at three logarithmically-spaced bin sizes (1, 1.6 and 3.5 ms) and TE was computed at delays (0-3 bins, for bins of size 1 and 1-4 bins for bins of size 1.6 and 3.5 ms) corresponding to synaptic delays, as in Timme et al. (2014, 2016). Thus, we computed TE at three timescales, 0.05–3 ms, 1.6–6.4 ms and 3.5–14 ms. Timescales were purposefully designed to be overlapping so that no interactions were neglected. See Supplemental Figure 1 for an overview of the binning structure used in TE calculations. To consider extra-synaptic interactions, we computed additional networks with bin sizes 7.5, 16.15, 34.8, 75, 161.6, 348.1, and 750 ms. With delays of 1-4 bins, these networks represent timescales of 7.5-30 ms, 16.15-64.6 ms, 34.8-139.2 ms, 75-300 ms, 161.6-646.6 ms, 348.1-1392.4 ms, and 750-3000 ms (Supplemental Table 1).

| **Timescale** | **Bin size (ms)** | **Delay window (ms)** | **Delay window (bins)** | **Timescale Delay (~ms)** |
| --- | --- | --- | --- | --- |
| 1 | 1 | 0.05–3 | 0–3 | 3 |
| 2 | 1.6 | 1.6–6.4 | 1–4 | 5 |
| 3 | 3.5 | 3.5–14 | 1–4 | 11 |
| 4 | 7.5 | 7.5–30 | 1–4 | 23 |
| 5 | 16.15 | 16.15–64.6 | 1–4 | 48 |
| 6 | 34.8 | 34.8–139.2 | 1–4 | 104 |
| 7 | 75 | 75–300 | 1–4 | 225 |
| 8 | 161.6 | 161.6–646.4 | 1–4 | 485 |
| 9 | 348.1 | 348.1–1392.4 | 1–4 | 1044 |
| 10 | 750 | 750–3000 | 1–4 | 2250 |

**Supplemental Table 1.** Timescales. Bin sizes and delays were chosen to logarithmically span a broad range of neurologically relevant time scales. Timescales were somewhat overlapping in order to ensure that all interactions were captured. In addition, timescales beyond 1 used delays in order to prevent short timescale interactions from influencing long timescale dynamics. Values in the far right column were used to refer to timescales throughout the paper in an abbreviated fashion. For additional information about the binning structure, see Supplemental Figure 1.

TE quantifies an effective connection from neuron J to neuron I by measuring how much information the past state of the neuron J time series ($J_{t-1}$ ) produces regarding the current state of the neuron I time series ($I_{t}$ *)*, beyond what is provided by the past state of the neuron I time series ($I_{t-1}$). Here, time series are binary spike trains for neurons I and J, containing 0 for time bins in which the neuron did not spike and 1 for time bins in which it did spike. Generally, the TE from neuron J to neuron I is computed as:

$${TE}_{J \to I}=\sum_{i_{t},i_{t-1}, j_{t-1}} p\left( i_{t}, i_{t-1}, j_{t-1} \right)\log\left( \frac{p(i_{t}| i_{t-1}, j_{t-1})}{p\left( i_{t} | i_{t-1} \right)} \right)$$

(1)

The probabilities in Eqn. 1 are computed by counting the number of occurrences of all possible combinations of spiking and not spiking in the $i_{t}, i_{t-1}$ and $j_{t-1}$ time bins (of the $I_{t}$, $I_{t-1}$ and $J_{t-1}$ time series) for all bins making up the hour-long recording.

Because we wanted to consider interactions at various timescales associated with synaptic and extra-synaptic transmission, we included a delay between the past and future states of the neurons so that $i_{t-1}$ became $i_{t-d}$ and $j_{t-1}$ became $j_{t-d}$. Additionally, in order to ensure overlapping timescales, we combined the $i_{t-d}$ and $j_{t-d}$ bins with their previous time bins, such that a spike in either or both time bins corresponded to a state of 1 while no spikes in either time bin corresponded to a state of 0 (see Supplemental Figure 1 for binning structure). Denoting these new bins as $i_{t-d}^{'}$ and $j_{t-d}^{'}$ gives a slightly different form for TE:

$${TE\left( d \right)}_{J \to I}=\sum_{i_{t},i_{t-d}^{'}, j_{t-d}^{'}} p\left( i_{t}, i_{t-d}^{'}, j_{t-d}^{'} \right)\log\left( \frac{p(i_{t}| i_{t-d}^{'}, j_{t-d}^{'})}{p\left( i_{t} | i_{t-d}^{'} \right)} \right)$$

(2)

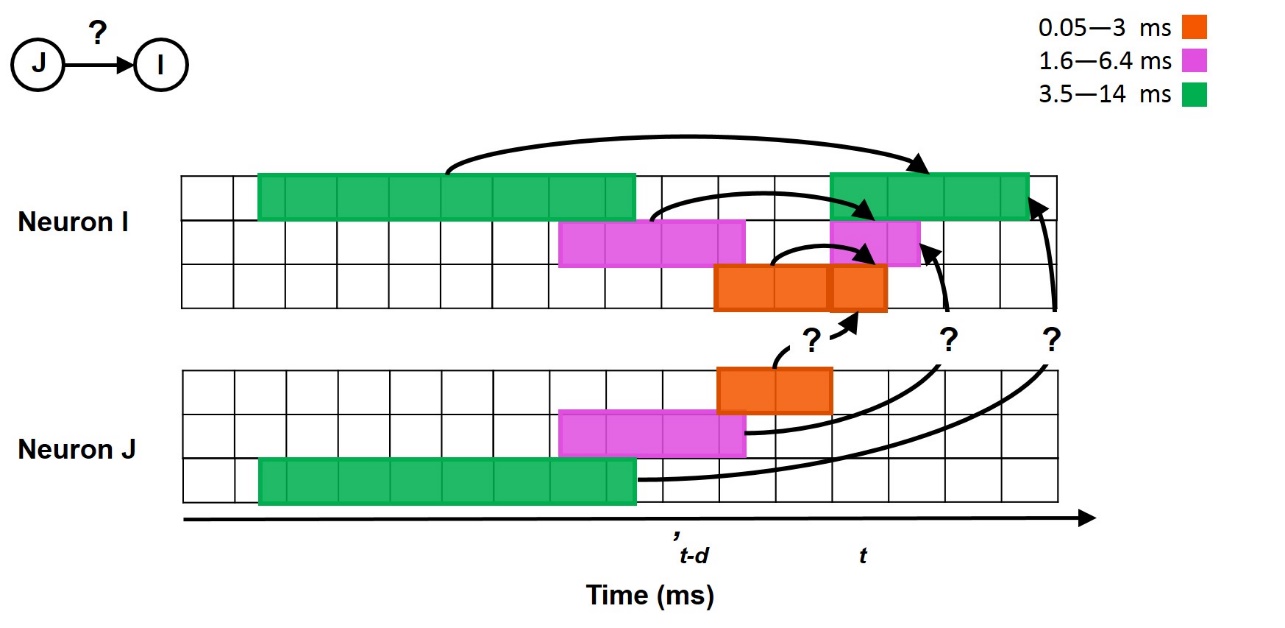

**Supplemental Figure 1.** Overview of time series binning structure used in transfer entropy calculations. Transfer entropy was used to quantify a directed, functional connection from neuron J to neuron I which represents how well the current state (t) of neuron I can be predicted by the past state (’t-d) of neuron J, beyond what is known from the past state of neuron I itself. Three synaptic timescales were considered, each with corresponding delays (d). These timescales considered transfer entropy from 0.05­—3 ms, 1.6—6.4 ms, and 3.5—14 ms.

To cast TE in terms of the percentage of the receiver neuron’s capacity that can be accounted for by the transmitting neuron, rather than it representing the amount of information being transmitted from transmitter to receiver, we normalized TE by the entropy of the receiver neuron via:

$${{TE}_{Norm}\left( d \right)}_{J \to I}= \frac{{TE\left( d \right)}_{J \to I}}{-\sum_{i_{t}} p(i_{t})log( p(i_{t}))}$$

(3)

Computing (normalized) TE in this way between all pairs of binned neuronal time series results in a time-scale dependent, weighted, directed network. Networks are weighted because some pairs of neurons fire more frequently and reliably at certain delays than others, and they are directed because a predictive, statistical relationship that exists from neuron J to neuron I, may not exist from neuron I to neuron J. Each element $a_{ij}$ in the TE matrix is the TE value from the $i^{th}$ to the $j^{th}$ neuron. TE values of zero denote the absence of an effective connection between the two neurons, while TE values greater than zero represent the weighted strength of the effective connection between the two neurons.

To determine the significance of network connections (TE values), TE values were computed for 5000 pairs of jittered spike trains. Spike trains were jittered by randomly adjusting the timing of each spike by a small amount proportional to the timescale being examined. This preserved the overall firing of each neuron, as well as the longer timescale dynamics, but destroyed the precise spike timing between neurons. Only spike trains of source neurons were jittered in order to preserve the auto-prediction in the target neuron. Spike trains were jittered according to a uniform distribution with a width of seven bins centered on the observed location of each spike. This procedure is discussed in more detail in Timme et al. 2014. TE values which were larger than 99.9% of jittered values were considered significant, corresponding to an alpha value of 0.001. Computing significant, normalized TE values for 25 recordings at ten timescales resulted in 250 full networks.

An important consideration in the calculation of our TE values, which includes a delay in the past of the receiver, is that we may be overestimating TE and increasing our odds for detecting false positives (Wibral et al., 2013). Importantly, we control for the increased risk of false-positives by using an alpha-level of 0.001. However, the potential overestimation of TE comes from more variance being attributed to the sender neurons than to the receiver neuron itself. Despite the advantages of the Wibral et al. method, which does not include a delay in the receiver past, we chose the current method in order to isolate interactions at separate timescales. It is possible that our choice to do so may have impacted the results of our non-normalized analysis of synergy with respect to mutual information (see Supplemental Results below). The non-normalized analysis resulted in ever-increasing information values across timescales, as delays lengthen. Yet, we do not observe this in our normalized results. Thus, the normalization of these values by the entropy of the receiver appears to prevent the delay confound from having a runaway effect. In addition, the normalization serves to enable direct comparison across timescales.

To be confident that our timescales captured the peak of information processing in our networks, we calculated TE at delays other those analyzed here for two representative networks. First, spike trains were binned at 1 ms. Then TE was calculated at multiple delays ranging from 0 to 501 ms, in steps of 3 ms, for all existing pairs of significant connections in the network. We found that TE tended to peak in the 1-14 ms delay range for most connections (Supplemental Figure 2). Across networks, 87.3% of pairs, on average, had a peak of TE between 1 and 14 ms.

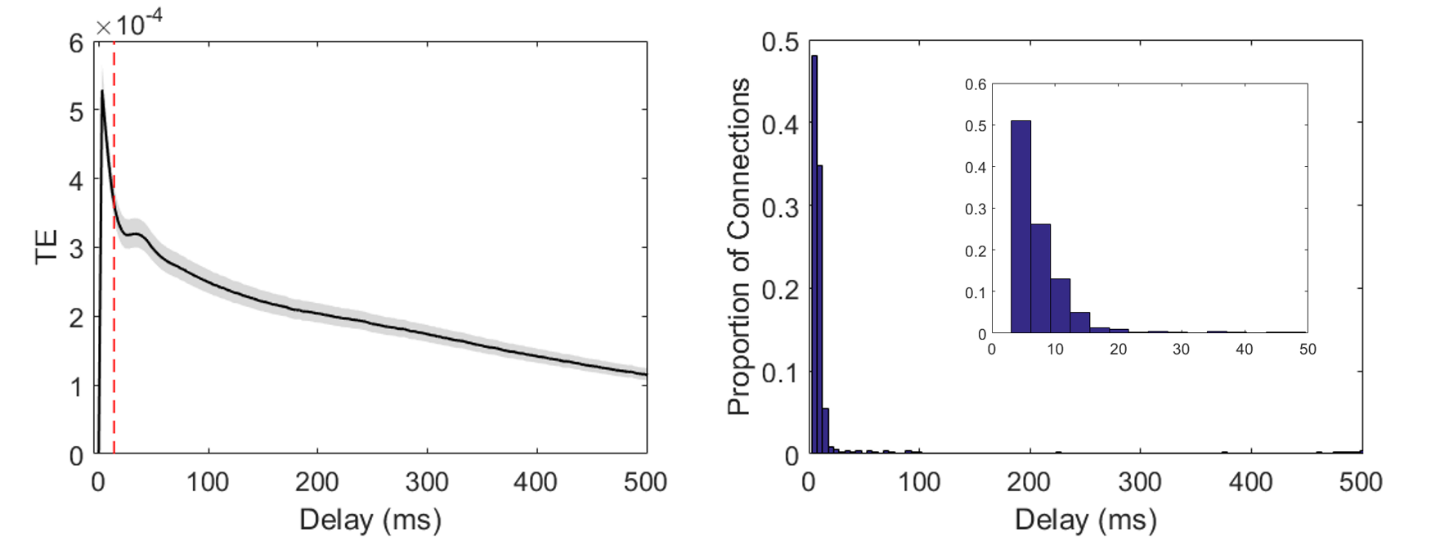

**Supplemental Figure 2.** TE peaks between 1-14 ms. Mean distribution of TE over time for all connections from two representative networks. Left: The black line shows the mean TE over all connections from two representative networks. The shaded region shows the 95% confidence interval. The vertical dashed red line indicates the upper bound of the synaptic timescales. Across connections, the peak TE occurs below this bound at short latencies. Right: Histogram of the delay to the maximum TE over connections. The height of each bar shows the proportion of connections for which the peak TE was found to occur at the delay indicated along the x-axis. Most connections had max TE at short delays as shown in the inset panel which zooms in to the first 50 ms of the x-axis. These plots show that most connections had a peak TE at less than 14 ms.

**Quantification of multivariate transfer entropy, redundancy, and synergy**

Computation by neurons receiving inputs from two other neurons in these networks was quantified following the partial information decomposition (PID) method from Williams and Beer (2011). The PID allows multivariate TE (mvTE) to be separated into distinct information components, one of which is a measure of neural computation termed synergy. The general form of the decomposition of multivariate TE between three neuronal time series, with two transmitter neurons, J and K, each sending a single input to one receiver neuron, I, can be expressed as:

$$TE\left( \left\{ J,K \right\}\to I \right)= \mathrm{Synergy}\left( \left\{ J,K \right\}\to I \right)+ \mathrm{Unique}\left( K;J\to I \right)+$$

$$\mathrm{Unique}\left( J;K\to I \right)+ \mathrm{Redundancy}(\{J,K\}\to I)$$

(4)

where $\left\{ J,K \right\}$ is a vector of the combined J and K time series (Supplemental Figure 3). Similarly, we can express the decomposition of bivariate TE from neuron J to I and neuron K to I as:

| $TE\left( J\to I \right)= \mathrm{Unique}\left( K;J\to I \right)+ \mathrm{Redundancy}(\{J,K\}\to I)$ | (5) |
| --- | --- |
| and |  |
| $TE\left( K\to I \right)= \mathrm{Unique}\left( J;K\to I \right)+ \mathrm{Redundancy}(\{J,K\}\to I)$ | (6) |

In Equations 4-6, all terms are quantified in units of bits (see Williams and Beer 2010, 2011 for a full description of these terms). The unique terms correspond to the information provided by that time series alone (either the J or the K time series) about the current state of I. The redundant term represents the overlapping information provided by time series J and K about the current state of I. Notice, in Equations 5 and 6, that although TE is only dependent on the two time series that are directly interacting (either J and I, or K and I), because J and K are both interacting with the same time series, their unique interactions are influenced by each other. Thus, the unique information provided by one of these time series is dependent on the other. In other words, because J and K provide some redundant (overlapping) information about I, J influences how much information K provides uniquely versus redundantly about I. Likewise, K influences how much information J provides uniquely versus redundantly about I.

The synergistic term in Equation 4 is the additional information (beyond the unique and redundant information) that is accounted for in the receiver activity (I) based on the non-overlapping information from both inputs (J and K) occurring simultaneously. Thus, synergy is a proxy for the non-linear computation which takes information from two sources and combines them in some way to generate a unique output. Examples of high synergy computations include ‘AND’ and ‘XOR,’ wherein knowledge of the state of both upstream neurons is needed to predict the state of the receiving neuron (Timme et al., 2016).

To calculate synergy, note that Equation 4 can be rewritten as:

$$\mathrm{Synergy}\left( \left\{ J,K \right\}\to I \right)= TE\left( \left\{ J,K \right\}\to I \right)- TE\left( J\to I \right)-$$

$$TE\left( K\to I \right)+ \mathrm{Redundancy}(\{J,K\}\to I)$$

(7)

by substituting Equations 5 and 6 and solving for synergy. Notice that we can compute all TE terms in Equation 7 via Equation 1. This leaves only the Redundancy term to be computed. We use the method for estimating redundancy provided by Williams and Beer (2010, 2011). That is redundancy is defined as follows in terms of a quantity titled the minimum information $I_{\min}$:

$$\mathrm{Redundancy}\left( \left\{ J,K \right\}\to I \right)≝I_{\min}\left( I_{t};J_{t-1}K_{t-1}|I_{t-1} \right)=$$

$$\sum_{i_{t}} p\left( i_{t} \right)\min_{R\in\left\{ J_{t-1},K_{t-1} \right\}} I_{spec}\left( I_{t}= i_{t};R | I_{t-1} \right)=$$

$$\sum_{i_{t}} p\left( i_{t} \right)\min_{R\in\left\{ J_{t-1},K_{t-1} \right\}} [I_{spec}\left( I_{t}=i_{t};R,I_{t-1} \right)-I_{spec}\left( I_{t}=i_{t};I_{t-1} \right)]$$

(8)

where the specific information $I_{spec}$ is defined as:

$$I_{spec}\left( I_{t}=i_{t};R,I_{t-1} \right)=\sum_{r, i_{t-1}} p\left( r,i_{t-1}|i_{t} \right)\log\left( \frac{p(r, i_{t-1}, i_{t})}{p\left( r,i_{t-1} \right)p(i_{t})} \right)$$

(9)

and

$$I_{spec}\left( I_{t}=i_{t};I_{t-1} \right)=\sum_{i_{t-1}} p\left( i_{t-1}|i_{t} \right)\log\left( \frac{p(i_{t-1}, i_{t})}{p\left( i_{t-1} \right)p(i_{t})} \right)$$

(10)

Thus, redundancy is the minimum information provided by J or K about each state of I, averaged over all possible states. In other words, redundancy is the minimum overlapping information (the shared information) that the past states of J and K provide about the current states of I. Redundancy was calculated via Equations 9 and 10. Finally, synergy was calculated via Equation 7. Computing synergy for all possible triads (for each neuron that received at least two inputs, all possible groupings of two input neurons and the receiver were considered) in all networks yields a single synergy value per triad. We then normalized synergy (as well as redundancy and mvTE) values by dividing by the entropy of the future state of I, as done in Equation 3.

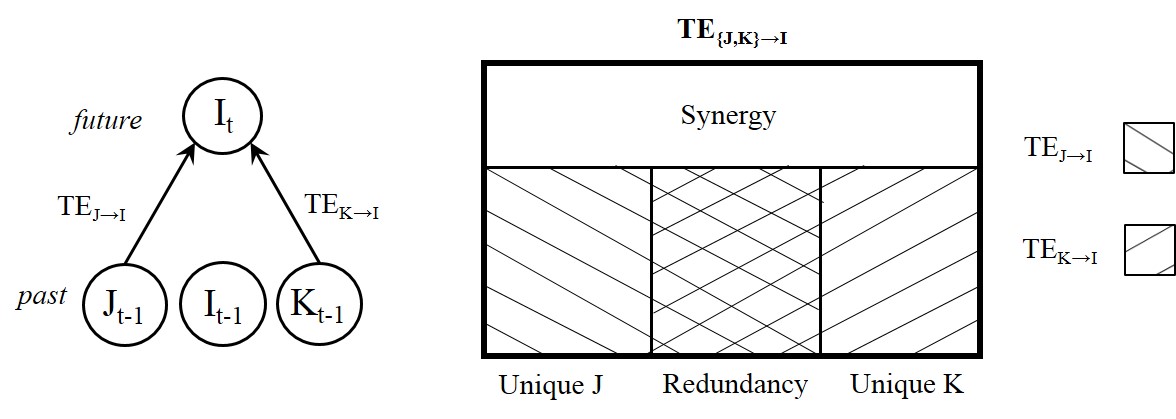

**Supplemental Figure 3.** The Partial Information Decomposition. In this study, we analyzed two-input computations which were determined using the Partial Information Decomposition to dissect multivariate transfer entropy (occurring among three neurons, with two transmitter neurons each sending significant information to a receiver neuron) into synergistic, redundant, and unique information terms. The synergistic information component was used to represent the amount of computation carried out by the receiver.

SUPPLEMENTAL RESULTS

Below are additional results not presented in the main text pertaining to (1) alternative measures of correlation (i.e., similarity) between senders; (2) non-normalized synergy versus mutual information; (3) an alternative implementation of synergy; and (4) linear regression slopes describing the synergy-mutual information relationship, as well relationships between synergy and redundancy and synergy and multivariate transfer entropy.

**Alternative Measures of Correlated Activity**

To test the robustness of the synergy-correlation relationship to analytical approach, we compared our results to those obtained from four alternative similarity metrics; these included: conditional mutual information, cosine similarity, Pearson correlation, and the Jaccard index (or Jaccard similarity). Their implementation and comparison to mutual information are described below, as are the results of their implementation. For each, the results are consistent with those obtained using mutual information as described in the main text.

Conditional mutual information is the mutual information between senders, conditioned on the past of the receiver. This alternative to mutual information was selected for its alignment to the multivariate TE calculation. Importantly, this metric removes variance related to the state of the receiver, thereby reducing the influence of the receiver on the mutual information between the senders. Cosine similarity computes the cosine of the angle between two spike trains, or the inner product of the spike trains, divided by the product of their magnitudes (e.g., Schreiber et al., 2003). This value scales between 0 and 1, with 0 being completely different (i.e., orthogonal) and 1 being identical. In comparison to mutual information, which is computed over all observations, cosine similarity ignores all observations of (0,0). Thus, this measure reflects the number of observations for which both neurons spike relative to spike rates of the two neurons. Pearson correlation coefficient computes the expectation of the variance of the two spike trains, divided by the product of their standard deviations. For binary spike trains, this is equivalent to the difference of the probability that one neuron spikes conditioned on whether the other spikes or does not. In comparison to mutual information, which weights each observation based upon the log-likelihood of observing that event type (i.e., (0,0), (0,1), (1,0), & (1,1) states), Pearson correlation weights each observation equally, independent of type. Values produced by this method scale between -1 and 1, with -1 being completely different, and 1 being identical. The Jaccard index computes the intersection of two spike trains, divided by their union. The Jaccard index scales between 0 and 1, with 0 being completely different, and 1 being identical. The Jaccard approach differs from the mutual information approach as it, like the cosine approach, ignores [0 0] observations. Unlike cosine approaches, it is not normalized by the firing rates of the two neurons, rather it is normalized by the total number of observations with at least one spike. For each of these alternate approaches to measuring the similarity of the transmitter neurons, we replaced the mutual information values with the respective alternate values and completed the remaining steps of the analyses identically.

The results show that the same qualitative pattern of findings emerge irrespective of which method is used for measuring the transmitter similarity. That is, regardless of the approach used to measure similarity, the median similarity increases as timescales grow longer, reflecting the transition from a low correlation regime to a high correlation regime as timescales shift from synaptic to extra-synaptic (Supplemental Figure 4). Also, regardless of the approach used to measure similarity there were positive correlations between similarity and synergy at synaptic timescales and there were negative correlations between similarity and synergy at extra-synaptic timescales (Supplemental Figure 5).

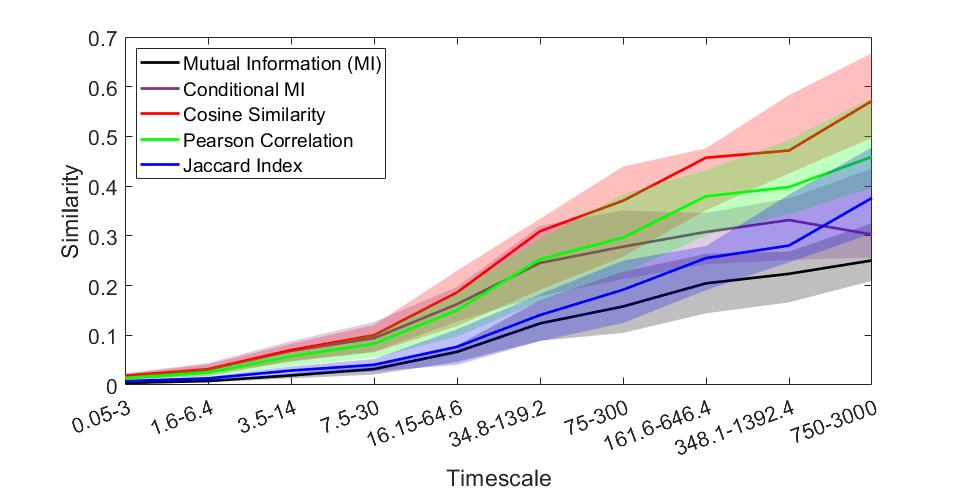

**Supplemental Figure 4**. Spiking activity of senders become increasingly similar at longer timescales irrespective of method for quantifying similarity. Lines indicate median median similarity across all recordings at each timescale. Shaded regions depict 95% bootstrap confidence intervals around the median.

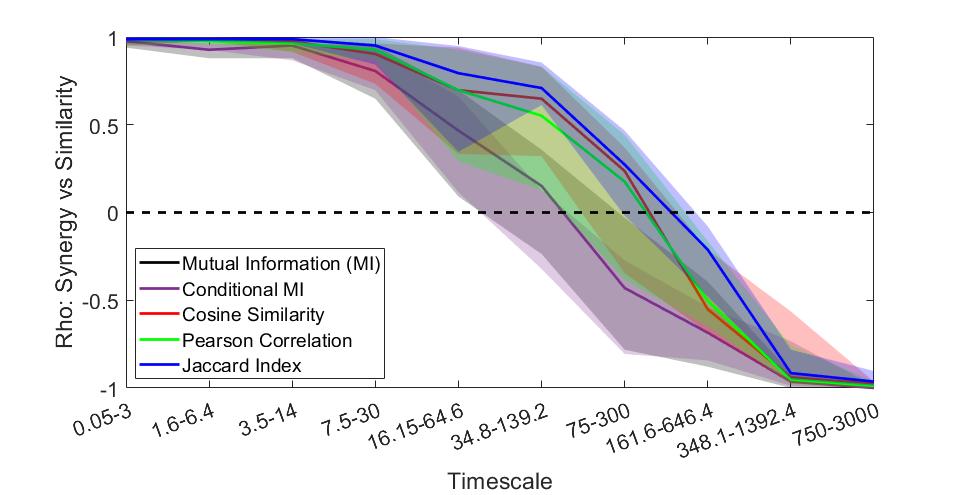

**Supplemental Figure 5**. Synergy is positively related to similarity at synaptic timescales (0.05-3 ms, 1.6-6.4 ms, and 3.5-14 ms) and negatively correlated with synergy at extra-synaptic timescales (75-300 ms, 161.6-646.4 ms, and 348.1-1392.4ms) irrespective of method for quantifying similarity. Lines indicate median correlation coefficients for similarity versus synergy for all recordings at each timescale. Shaded regions depict 95% bootstrap confidence intervals around the median. Conditional MI (purple) and MI (black) lines are overlapping here.

**Non-normalized synergy versus mutual information**

The results presented in the main text were obtained using mutual information and synergy values that were normalized by the maximum possible values given the firing rates and bin sizes for each triad at each timescale. This was done because information terms such as mutual information and synergy are sensitive to the entropy of the data being analyzed, which is, in turn, sensitive to firing rates and bin sizes. The log-normal distribution of firing rates and the different timescales generated large variability in the total entropy and thus information terms. By normalizing, we minimized any effects that may have been due to changes in entropy within and across timescales. To confirm that this did not influence our findings, we repeated the main analyses with no normalization. Supplemental Figure 6 illustrates the results showing that the same qualitative pattern of results with the non-normalized values.

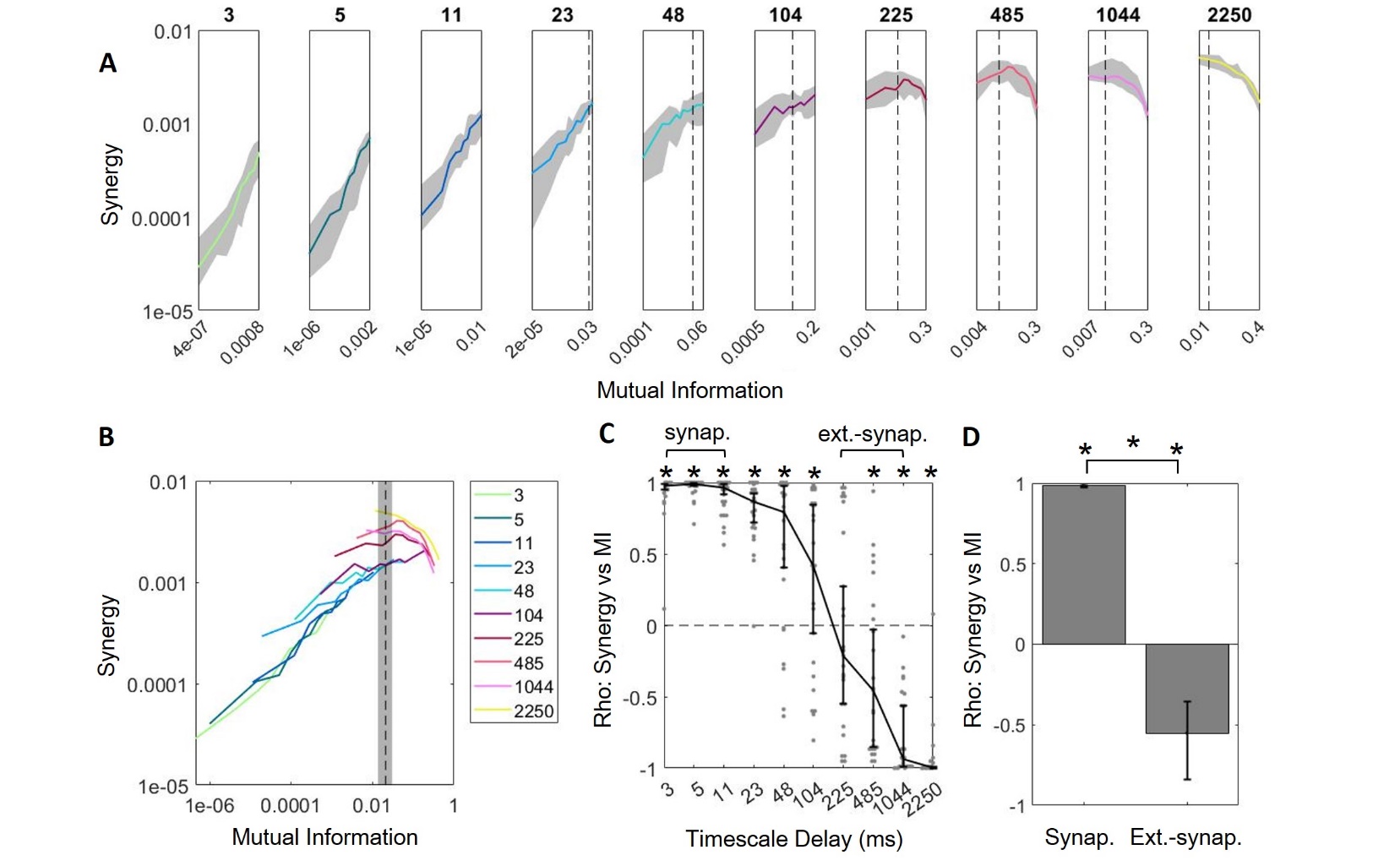

**Supplemental Figure 6.** Non-normalized synergy and mutual information values show the same qualitative trends as normalized values. (A) Plotting synergy as a function of mutual information across timescales shows that the positive relationship between synergy and mutual information only exists at peri-synaptic timescales where mutual information is relatively low. This relationship becomes negative at longer timescales where mutual information is high. Note that mutual information, plotted along the x-axis, varies across sub-panels. A dashed vertical line where mutual information is 0.02 is included to facilitate visual alignment across panels. (B) Curves from A are replotted on the same axes here to show the overall relationship between synergy and mutual information. (C) Spearman rank correlation coefficients of synergy versus mutual information for all networks across timescales show that synergy and mutual information are positively related at shorter timescales and negatively related at longer timescales. Correlation coefficients are largest for synaptic timescales. (D) Synergy and mutual information are significantly positively correlated at synaptic timescales and significantly negatively correlated at extra-synaptic timescales. For all plots, solid/bold lines indicate medians and shaded regions/error bars depict 95% bootstrap confidence intervals around the median.

**Alternative implementation of synergy**

To ensure that our findings were not dependent on the method used to calculate synergy, we performed additional analyses which implemented an alternative method. In this alternative method, we considered the effect of calculating the lower bound on synergy, which we refer to as “bonafide” synergy. This method also uses PID but sets redundancy to be the smallest possible value. Effectively, in this approach synergy is computed as follows:

$$\mathrm{Synergy}\left( \left\{ J,K \right\}\to I \right)= \mathrm{argMax}[ TE(\{J,K\}\to I) - TE\left( J\to I \right) - TE\left( K\to I \right), 0 ]$$

(11)

Consequently, synergy is minimized or set to zero when the sum of $TE(J\to I)$ and $TE(K\to I$) is greater than$TE(\{J,K\}\to I)$. To avoid having the number of zeros in any decile driving the results, the analyses were restricted to triads with synergy greater than zero.

When we used these synergy values and repeated our core analyses, we found that synergy and mutual information were positively related at synaptic timescales. However, we also found them to be positively related at extra-synaptic timescales, departing from what we show in the main text (Supplemental Figure 7). This is likely due to the fact that, in triads with sufficiently high synergy to be included in this analysis, redundancy had not effectively cannibalized the mvTE. As a result, the only triads left in this analysis were those with generally high synergy.

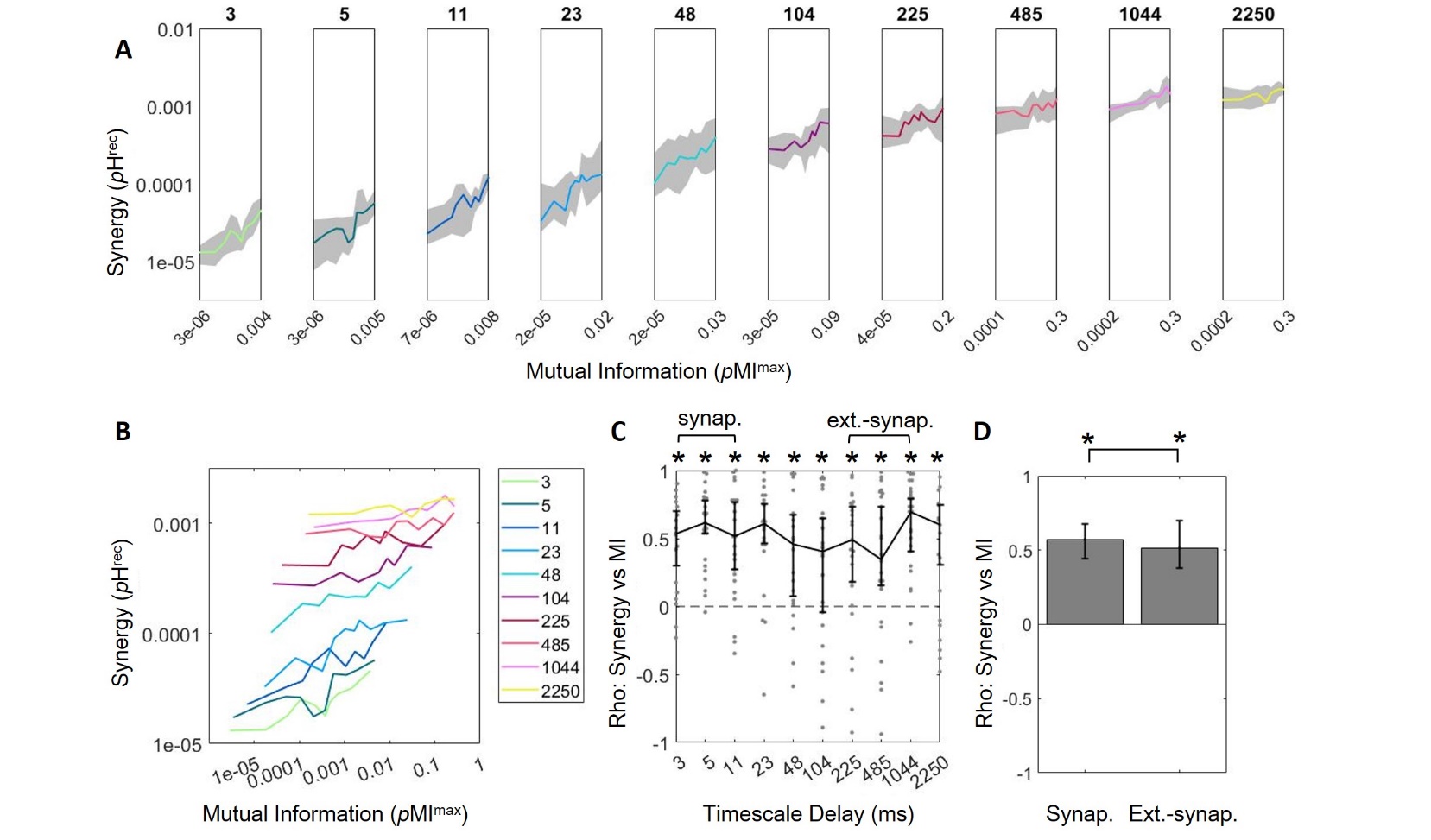

**Supplemental Figure 7**. Bonafide synergy is positively related to Sender_MI_ at all timescales. (A) Log-scaled Synergy versus mutual information grows at all timescales. (B) Curves of synergy versus mutual information in A, replotted on the same axes. All curves are increasing. (C) Spearman rank correlation coefficients for synergy versus mutual information at all timescales show that the two are always positively related. (D) Synergy and mutual information are positively related at both synaptic and extra-synaptic timescales. For all plots, solid/bold lines indicate medians and shaded regions/error bars depict 95% bootstrap confidence intervals around the median.

**Linear regression slopes of information term relationships**

In the main text we show Spearman rank correlation coefficients for synergy versus mutual information across networks (main text Figure 4), as well as other information term relationships (main text Figure 5). However, we also performed linear regressions in order to measure the slopes of these relationships. These are shown below in Supplemental Figure 8.

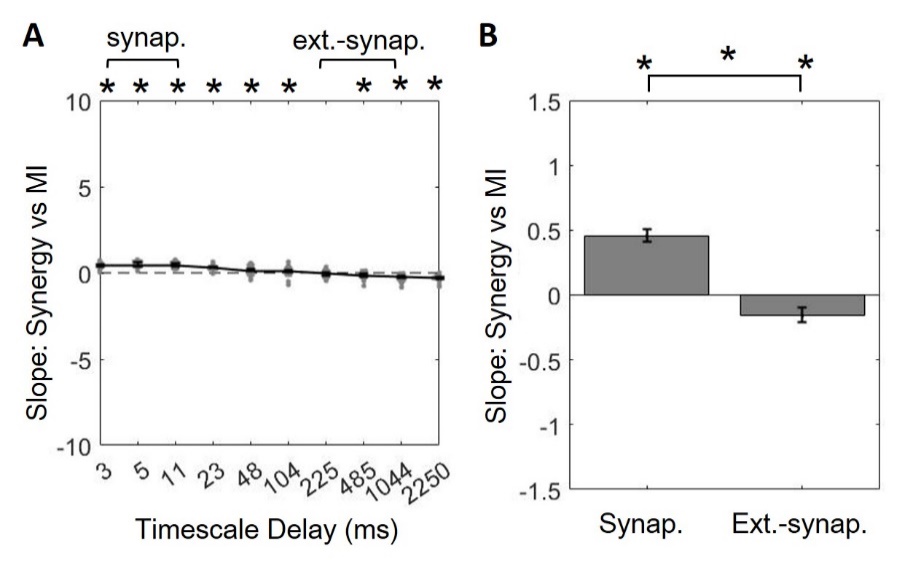

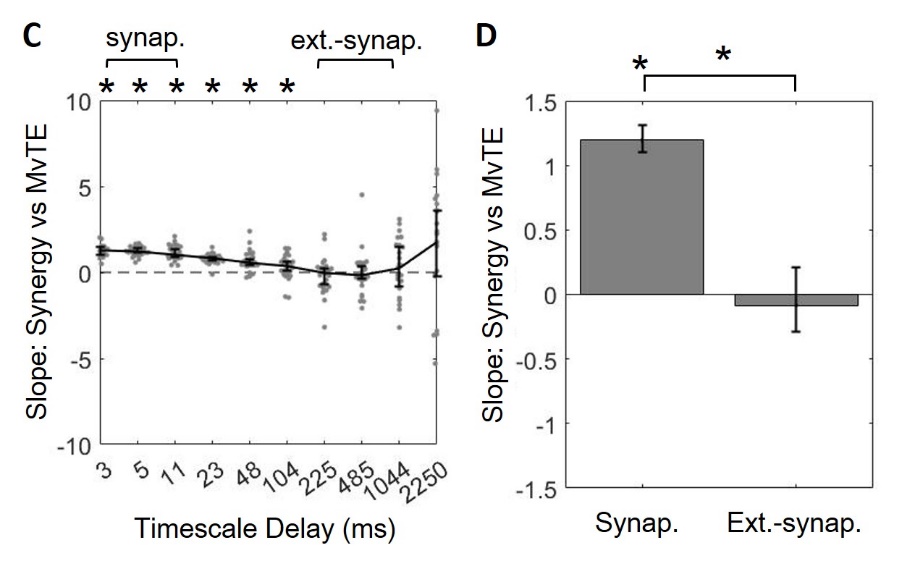

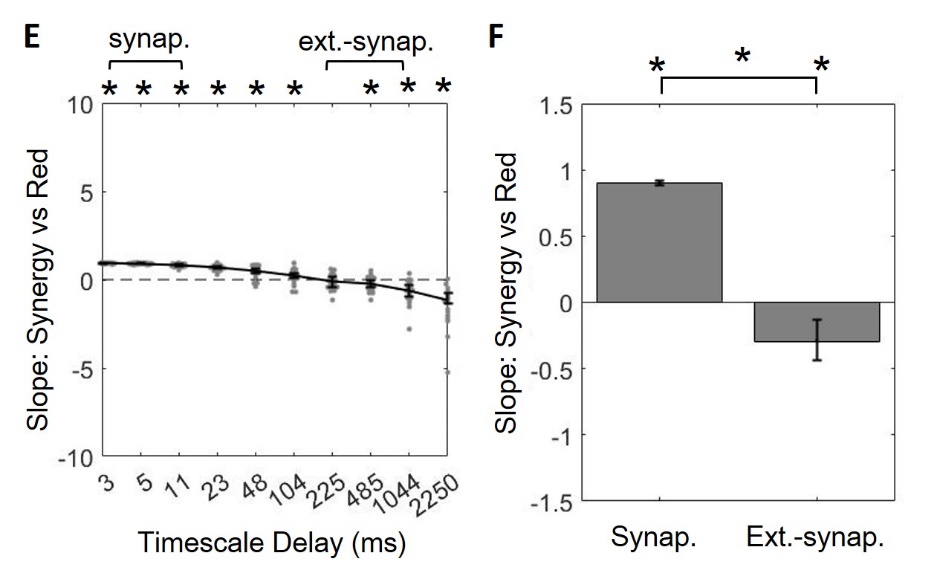

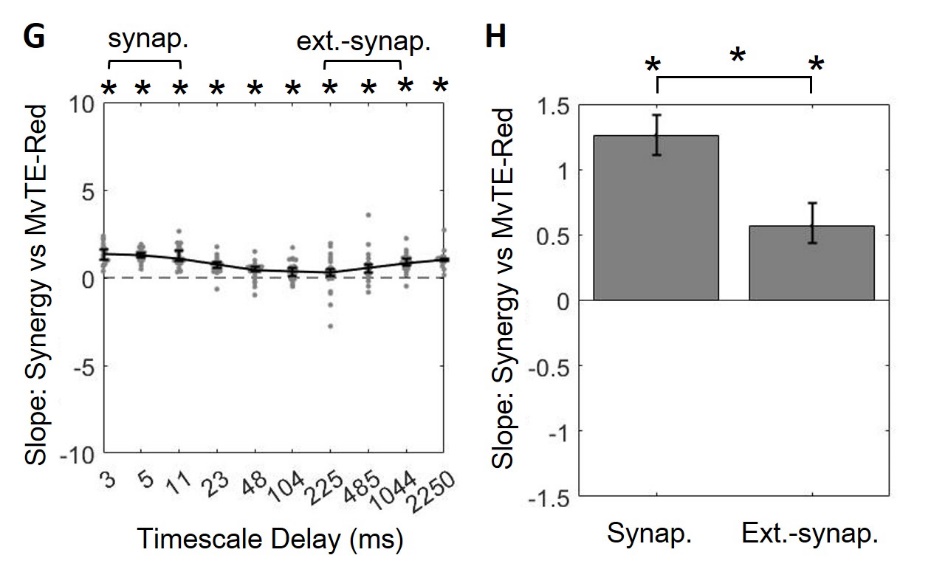

**Supplemental Figure 8.** Slopes of information term relationships show the same trends as Spearman rank correlation coefficients. (A) Synergy and mutual information are initially positively related, but become negatively related across timescales [ Δlog_10_(*p*MI_max_) / Δlog_10_(*p*H_rec_) ]. (B) Synergy and mutual information are positively related at synaptic timescales and negatively related at extra-synaptic timescales. (C) Synergy and mvTE are initially positively related, but become increasingly unrelated across timescales [ Δlog_10_(*p*H_rec_) / Δlog_10_(*p*H_rec_) ]. (D) Synergy and mvTE are positively related at synaptic timescales and unrelated related at extra-synaptic timescales. (E) Synergy and redundancy are initially positively related, but become negatively related across timescales [ Δlog_10_(*p*H_rec_) / Δlog_10_(*p*H_rec_) ]. (F) Synergy and redundancy are positively related at synaptic timescales and negatively related at extra-synaptic timescales. (G) Synergy and the difference between mvTE and redundancy are positively related at all timescales [ Δlog_10_(*p*H_rec_) / Δlog_10_(*p*H_rec_) ]. (H) Synergy and mvTE-redundancy are positively related at synaptic timescales and at extra-synaptic timescales. For all plots, solid/bold lines indicate medians and shaded regions/error bars depict 95% bootstrap confidence intervals around the median.
